# Immunological and cardio-vascular pathologies associated with SARS-CoV-2 infection in golden syrian hamster

**DOI:** 10.1101/2021.01.11.426080

**Authors:** Zaigham Abbas Rizvi, Rajdeep Dalal, Srikant Sadhu, Yashwant Kumar, Tripti Shrivastava, Sonu Kumar Gupta, Suruchi Agarwal, Manas Ranjan Tripathy, Amit Kumar Yadav, Guruprasad R. Medigeshi, Amit Kumar Pandey, Sweety Samal, Shailendra Asthana, Amit Awasthi

## Abstract

Severe acute respiratory syndrome coronavirus (SARS-CoV)-2 infection in golden Syrian hamster (GSH) causes lung pathology and resembles human coronavirus disease (Covid-19). However, extra-pulmonary pathologies of SARS-CoV-2 infection that result in long Covid remains undefined in GSH. Here, using *in silico* modelling we show that hamster angiotensin-converting enzyme-2 (ACE-2) and neuropilin-1 (NRP-1) interaction with SARS-CoV-2 is similar to human. Intranasal SARS-CoV-2 infection in GSH resulted in early onset of lung pathologies marked by aggressive inflammatory response. Remarkably, late phase of SARS-CoV2 infection in GSH showed cardiovascular complications (CVC) characterized by ventricular hypertrophy, ventricular wall thickening, interstitial coronary fibrosis and altered lipidomics with elevated cholesterol, low-density lipoprotein and long chain fatty acid triglycerides. Moreover, serum metabolomics profile of infected GSH correlated with Covid19 patients. Together, we propose GSH as a suitable animal model to study immediate and long Covid19 pathologies that could be extended to therapeutics against Covid19 related CVC.

## Introduction

First reported in Wuhan, China in December 2019, SARS-CoV-2 has infected nearly 0.9 % of the total world population with around 2.17 % mortality rate as of January 2021 (https://covid19.who.int/). Symptomatic COVID-19 is typically characterized by symptoms ranging from mild to acute respiratory distress which is associated with cytokine storm (Chen and Li, 2020; Verity et al., 2020). Moreover, extra-pulmonary symptoms ranging from CVC, inflammation, coagulopathies, multiple organ damage and neurological disorders that are also recognized as long term consequences of Covid19 or Long Covid have been described in patients with mild to severe covid-19 (Chen and Li, 2020; Cortinovis et al., 2021; Huang et al., 2021; Lamers et al., 2020; Mao et al., 2020; Neurath, 2020; Nishiga et al., 2020; Y. Wu et al., 2020; Xiao et al., 2020; Xydakis et al., 2020). The severity of covid-19 is governed by several host-virus factors such as dose of virus, route of viral entry, gender, age and comorbidity (Chen and Li, 2020; Sungnak et al., 2020; Wang et al., 2020). The RBD of spike (S) glycoprotein of SARS-CoV2 engages with angiotensin-converting enzyme 2 (ACE2), a cellular receptor expressed on host cells, which facilitates the viral entry into the host cell. Upon the engagement of ACE2 with SARS-CoV2, cellular transmembrane protease ‘serine 2’ (TMPRSS2) mediates the priming of viral S protein by cleaving at S1/S2 site induces the fusion of viral and host cellular membranes, thus facilitating viral entry into the cells (Hoffmann et al., 2020). Lower levels of surface expression of ACE2 in respiratory and olfactory epithelial cells, however, indicates a possibility of involvement of other co-receptors, which may be crucial for the infectivity of SARS-CoV2. Neuropilin-1 (NRP-1), which is abundantly expressed on the surface of endothelial and epithelial cells and binds to furin cleaved substrates, has been shown to facilitate SARS-CoV2 infectivity (Cantuti-Castelvetri et al., 2020; Daly et al., 2020). S1 fragment of the spike protein cleaved by furin has been shown to directly bind to NRP1 expressed on the cell surface, and thus promote the viral interaction with ACE2 (Cantuti-Castelvetri et al., 2020; Daly et al., 2020).

Various non-human primates and small animal models have been described to study the pathogenesis and transmission of SARS-CoV2 infection(Callaway, 2020; CohenApr. 13 et al., 2020; Johansen et al., 2020; Lakdawala and Menachery, 2020; Shi et al., 2020). SARS-CoV2 is inefficient in infecting mice due to structural differences in mouse ACE2 as compared to human ACE2, thus wild type mice are not the natural host to study SARS-CoV2 induced pathologies (Bao et al., 2020, p. 2; Chan et al., 2020). On the other hand, GSH, which was previously described as a model for SARS-CoV infection, has recently gained much attention as a suitable model for studying SARS-CoV2 infection (Chan et al., 2020; Imai et al., 2020; Kaptein et al., 2020; Kreye et al., 2020; Lee et al., 2020; Osterrieder et al., 2020; Sia et al., 2020; Tostanoski et al., 2020). Remarkably, hamsters have been shown to be infected through intranasal, oral and ophthalmic routes by SARS-CoV2 with respectively descending lung viral load and pathologies (Imai et al., 2020; Lee et al., 2020). Despite several reports on hamsters describing the SARS-CoV2 pathology in the lung and pre-clinical evaluation of therapeutics against it, no study has focused on extra-pulmonary pathologies that are associated with long-term consequences of SARS-CoV2 infection.

In the current study, we demonstrate using *in-silico* modeling, that host-virus interaction has striking similarity between humans and hamsters that involves TMPRSS2-mediated priming of S protein, binding of S1 peptide with NRP-1 and subsequent binding of RBD to ACE2. Intranasal infection of SARS-CoV2 in hamster resulted in high lung viral load with significantly elevated lung injury on 2 and 4 days post-infection (dpi). Furthermore, our data shows aggressive immune activation characterized by heightened expression of inflammatory cytokines on 2 and 4 dpi. Strikingly, SARS-CoV2 infection in the hamster leads to significant interstitial coronary fibrosis on 7 and 14 dpi characterized by thickening of ventricular walls and interventricular septum. These cardiovascular changes were associated with increased serum triglycerides, low-density lipoprotein (LDL), cholesterol, high-density lipoprotein (HDL) on 4 dpi, and a marked increase in long-chain fatty acids (LCFAs) on 7dpi. Finally, we found marked changes in the metabolomics profile characterized by elevated N-acetylneuraminate and allantoin, which corroborated with the covid19 severe patients’ metabolomic profile. Taken together, here we provide evidence that SARS-CoV2 infection in GSH results in long Covid marked by CVC long after viral clearance from the lungs. We thus propose hamster as an excellent model to study both immediate as well as long-term consequences of SARS-CoV2 infection.

## Results

### *In silico* interaction predicts similarities in the host-virus interaction between hamster and human

The respective interactions of SARS-CoV-2 RBD and a fragment of S1 protein with host ACE2 receptor and NRP-1 are critical for virus entry into epithelial cells (Chan et al., 2020; Hoffmann et al., 2020). Since GSH mimics human SARS-CoV-2 infection, therefore it is essential to understand the similarities between human (hu) and GSH (ha) ACE2 receptor, TMPRSS2 and NRP-1 at the molecular level. To compare amino acid residues of hu and ha ACE2 that interact with RBD of SARS-CoV2, structure-guided sequence alignment was performed with their respective sequences (**Figures S1A and 1A**). The overlay of the crystal of huACE2 with modeled haACE2 was 1.2 Å, indicating the high structural resemblance. At the interface, 20 residues of hu ACE2 interact with 17 residues of RBD while 20 residues of ha ACE2 contact with 16 residues of RBD **(Figures 1B, IC, ID and IE).** The notable features that were common for both hu/ha ACE2 interfaces with RBD were the networks of hydrophilic interactions like hydrogen bonds (H-bonds) and salt bridges between Lys31 and Lys353 of ACE2 with Glu35 and Asp38 of RBD **(Figures 1E, 1F)**. Aromatic residues of RBD, Tyr449, Tyr453, Phe456, Phe486, Tyr489, Tyr495 and Tyr505, contributing to binding affinity remained common for both hu/ha ACE2 **(Figure 1F)**. Together, SARS-CoV2 RBD with hu/ha ACE2 interfaces shared substantial similarities in the buried surface area, the number of interacting residues and hydrophilic interaction. hu/ha TMPRSS2 structurally looks very similar (**Figure S1B)**. NRP-1, which binds to S1 fragment of spike protein, facilitates SARS-CoV2 entry within the cells (Cantuti-Castelvetri et al., 2020; Daly et al., 2020). Hu and ha NRP-1, the C-terminal R685 of the CendR peptide, interacts identically with S1 fragment of spike protein residues, Y297, W301, T316, D320, S346, T349 and Y353 (**Figures 1G, 1H and 1I**). The R682 and R685 sidechains S1 fragment of spike protein engages with NRP-1 sidechains of Y297 and Y353 via stacked cation-π interactions (**Figures 1I and 1J**). In huNRP-1, the residues interacting from C-endR are R685 making H-bonds with residues Asp320, Ile415, Tyr353, Thr349 and Ser346 of S1 fragment while in haNRP-1, the R685 remains stable but with less number of H-bonds with residues Trp297, Gly318, Asp320, Tyr353 and Thr349. The other common interactions in terms of hydrophobic contacts are with residues Thr316, Gly414, and Trp301. Glu348 and Lys351 contributed only in huNRP-1 while Ser346 in haNRP-1. Among mutations at RBD, the cause of stable Y501 over N501 was also compared in hu/ha, as Y501 gain additional bonding with tyrosine residues (41@ACE2, 355@RBD) and H-bond with Arg353 (**Figure 1K)**. Based on these molecular interaction analyses, it is suggested that hu and ha host proteins show similarities in their interactions with SARS-CoV2 proteins and further provided the basis of rational drug designing to block SARS-CoV2 entry within the cells.

**Figure 1.**
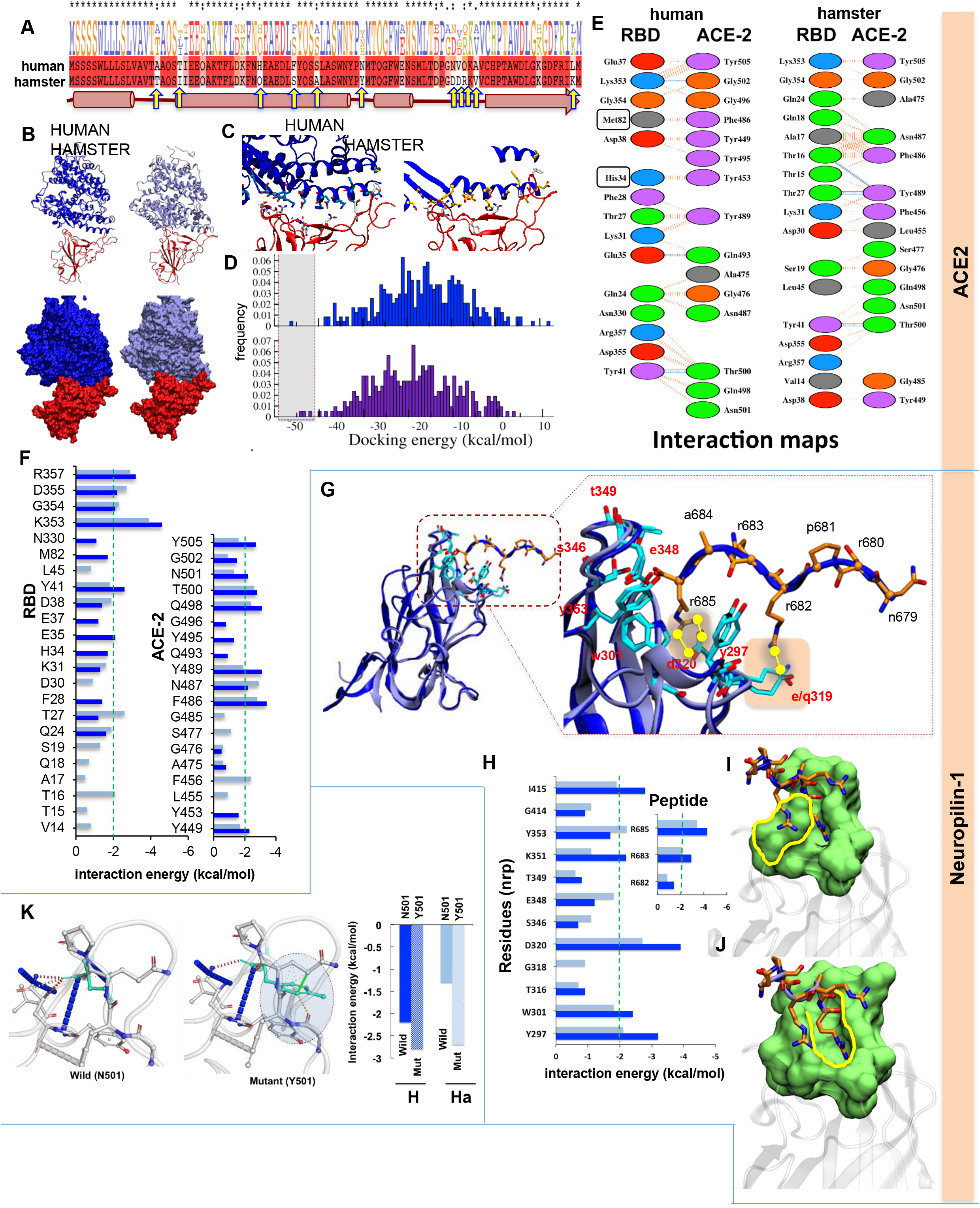
Computational analysis deciphering the structural and interaction level insights of key host proteins, ACE2 (A to F), NRP-1 (G to J) and N501Y (K) involved in SARS-CoV-2 infection. **(A)** the sequence alignment of human and hamster ACE2 sequences. Along with secondary structure, the residual variations are highlighted in yellow arrows indicating the differences among sequences. **(B)** crystal structure of human ACE2. The hACE2 (in blue), modeled haACE2 (in ice-blue), and RBD is in red. The surface view of h/ha ACE2 with RBD. **(C)** The residue level interaction map, ACE2, is rendered in cartoon and colored for h(blue) and for hamster (ice-blue). The amino acid residues are coloured as per atom-type such as O:red, N:blue, C:cyan/yellow/white. **(D)** The molecular docking data of ACE2 and RBD interaction (in Histogram). The selective docked pose is shown in a dotted rectangular box. **(E and F)** the interaction energies calculated at h/ha interfaces. The interacting residues are aromatic:purple, polar:green, acidic:red and aliphatic:grey. The types of interaction below <−3.0 Å distance are shown by blue lines and hydrophobic interactions are in orange dotted lines. The bar graphs are the residue-wise interaction energies (in kcal/mol) for both RBD and ACE2. The key residues (<−2.0 kcal/mol) are highlighted via green dotted lines. **(G)** The overlay of h/ha NRP-1 along with CendR peptide, rendered in cartoon. The zoom-out view of interacting interfaces of h/ha NRP-1 loop residues and CendR peptide residues. Peptide residues are rendered in licorice and color-coded in atom-wise. The hydrogen bond-forming residues are shown by yellow lines. **(H)** the bar graph showing the residue-wise interaction energy. **(I and J)** the binding groove of NRP-1 showing how nicely CendR peptides fit in h/ha. **(K)** The N501 and Y501 interaction patterns in hACE2-RBD. The bar graph showing the difference of 501 in h/ha in wild and mutant states.

### SARS-CoV2 infection results in pulmonary pathologies characterized by inflammation and lung injury

The *in silico* data provides a comparative molecular insight of SARS-CoV2 interaction and entry in GSH showing similarities with human infection. Further, a decrease in body mass had been reported in the initial phase of SARS-CoV2 infection. Inline, intranasal SARS-CoV2 infection in GSH showed a gradual body mass loss peaking (~10% as compared to uninfected) at 4 dpi with 10^4^ and 10^5^ plaque-forming unit (PFU) (**Figure 2A**) (Chan et al., 2020; Imai et al., 2020; Lee et al., 2020; Sia et al., 2020). This was accompanied by gross lung morphological changes characterized by pneumonitis regions with a high viral dose 10^5^ (PFU). TCID_50_ was found ~10.5 log_10_ copy number/g (lung mass) and at 2 dpi that gradually decreased at subsequent time points, marking the recovery of the animals from SARS-CoV2 infection (**Figures 2B and 2C)**. Covid-19 is marked by severity of pneumonia, lung injury and an influx of activated immune cells (Afrin et al., 2020; Chen and Li, 2020; Moore and June, 2020). In line with this, histological analysis revealed an elevated disease index score characterized by scores of inflammation, epithelial injury, lung injury and pneumonitis with elevated infiltration of granulocytes and mast cells on 2, 4 and 7 dpi (**Figures 2D and 2E**). Consistently, mRNA expression of mast cells signature enzymes, chymase and tryptase, was increased on 2 and 4 dpi corroborating mast cell enrichment (**Figure 2F**). Furthermore, the expression of eotaxin, a CC chemokine and a potent chemoattractant for airway eosinophils and mast cells that is particularly elevated in asthma and allergy conditions, was upregulated at 2 dpi (**Figure 2G**) (Guo et al., 2001). The expression of other lung injury markers like mucin (muc-1: marker of respiratory infections), surfactant protein-D (sftp-D: acute lung injury marker), advanced glycation end product (AGER: pro-inflammatory pattern recognition receptor) and plasmonigen activator inhibitor-I (PAI-1: a key factor for lung fibrosis) was upregulated at 4 dpi (**Figure 2G**) (Chatterjee et al., 2020; Crouch, 2000; Oczypok et al., 2017; Prabhakaran et al., 2003). Collectively, our data show that SARS-CoV2 infection in GSH results in high viral load at the onset of infection that is characterized by lung injury and inflammation.

**Figure 2.**
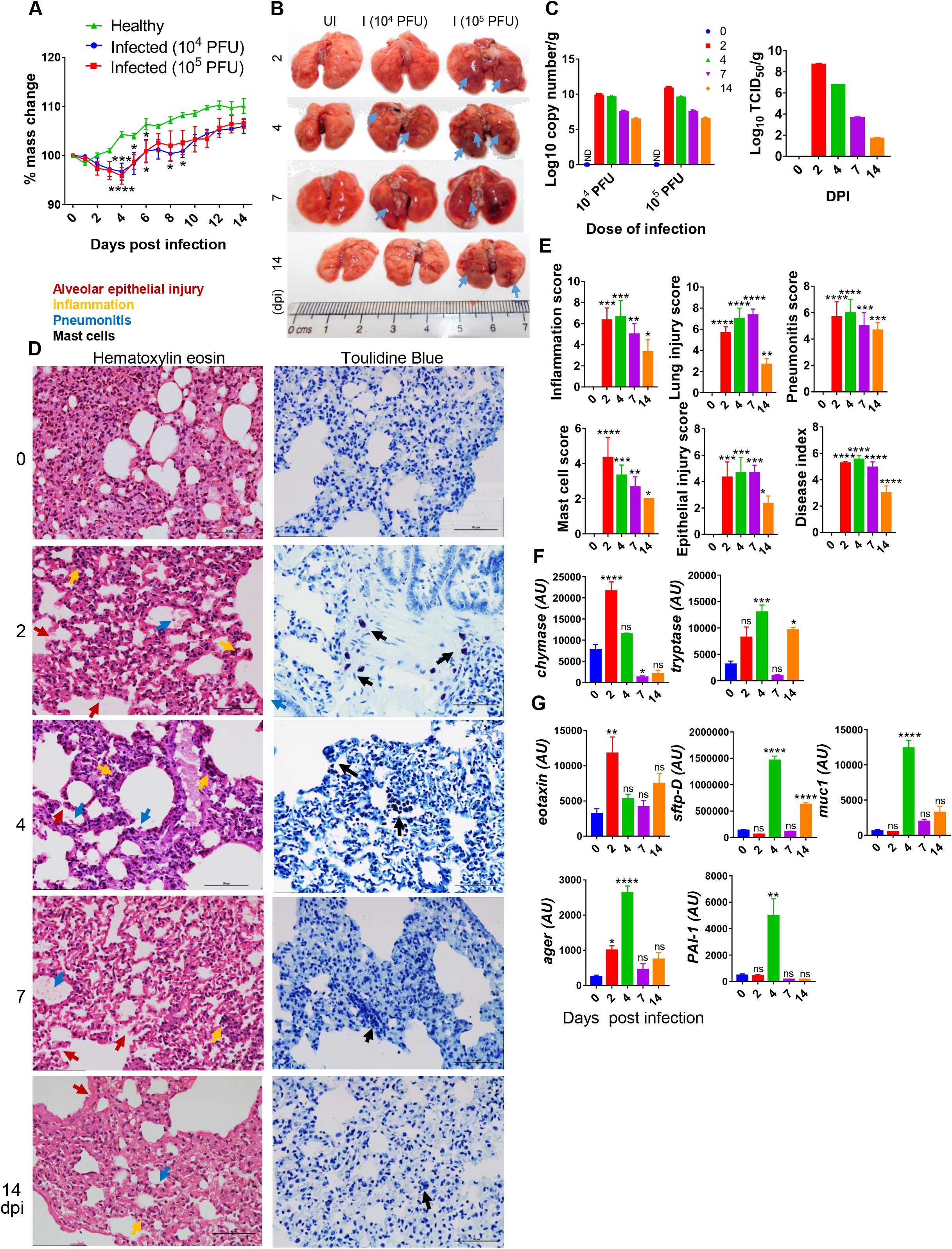
Pulmonary pathologies of SARS-CoV2 infected hamsters. (A) % body mass change. (B) Gross morphology of lungs showing pneumonitis region. (C) Lung viral load expressed as Log10 copy number/g or log10 TCID50/g. (D) Microscopic images of HE and TB stained lungs (E) histological scores (F & G) lung qPCR showing mean relative mRNA expression with standard error mean (SEM). Alveolar epithelial injury (red), inflammation (blue), pneumonitis (yellow) and mast cells (black). *P < 0.05, **P < 0.01, ***P < 0.001, ****P < 0.0001 (one-way anova).

### SARS-CoV2 infection in hamster causes early and aggressive onset of immune response

Most SARS-CoV2 infection studies in hamsters have demonstrated the pathological changes associated with the dissemination of virus in various organs without characterization of an acute immune response. Further, pathological changes of the lung and other organs have been attributed to the acute cytokine storm caused by SARS-CoV2 infection as shown in Covid19 patients (Iwasaki and Yang, 2020; Mathew et al., 2020; Moore and June, 2020; Verity et al., 2020). In line with this, profound splenomegaly was observed in infected animals with up to ~60% increase in spleen size on 4 dpi with 10^5^ PFU SARS-CoV2 infected GSH as compared to the healthy control (**Figure 3A**). These observations demonstrated that splenomegaly could be an indicator of disease severity to identify the efficacy of therapeutics against SARS-CoV2 infection in GSH. Splenomegaly was subsided at subsequent time points (7 and 14 dpi), which corroborate with recovery in the lung pathologies (**Figure 3A)**. To further corroborate with the recovery response of SARS-CoV2 infection in GSH, serum IgG titre for viral proteins, RBD, Spike and N protein, was determined. As compared to low dose, high dose of infection generate higher IgG titre against RBD (**Figure S2A**). The RBD, Spike (prefusion S2P), and N protein ELISA detected the corresponding antibodies in challenged animals as early as 7 dpi. The serum endpoint antibody titres against RBD, spike and N protein were detected as early as 7 dpi with the titre 1:3610, 1:405 and 1:330 respectively, the titre of these antibodies was further enhanced to 1:8900, 1:550 and 1:660 at 14 dpi (**Figure 3B**), indicating for a steady recovery of GSH from SARS-CoV2 infection. The cellular response was corroborated with antibody response, as the frequency of RBD-specific CD4^+^IFNγ^+^ cells was also found to be elevated (~2.5 fold) (**Figure 3C)**. The frequency of CD4^+^ T cells remained the same at 2 dpi between uninfected and SARS-CoV2 infected GSH (**Figure S2B**). To define cellular response, particularly T helper (Th) cell response, against SARS-CoV2 infection in GSH, we performed mRNA expression of crucial cytokines, chemokines, transcription factors and checkpoint inhibitors. Our data demonstrated an increased expression of signature cytokines of Th1 (IFNγ, tumor necrosis factor (TNF)-α), Th2 (interleukin (IL)-13), Th9 (IL-9) and Th17 (IL-17A) cells at the peak of infection (i.e. 2 dpi) along with their respective transcription factors, T-bet, GATA3, RAR-related orphan receptor (ROR)-c (Candia et al., 2021; Grifoni et al., 2020; Mathew et al., 2020). However, transcription factor forkhead box protein (Foxp)-3 was found to be elevated at 14 dpi, suggesting the activation of regulatory T (Tregs) cells at the senescence of infection. Interestingly, Tregs cytokines IL-10 and transforming growth factor (TGF)-β sets in much early at 2 dpi which maybe to counterbalance aggressive inflammatory reaction (**Figures 3D and 3E)**. We also found elevated expression of IL-6 and inducible nitric oxide synthase (inos) at 2 dpi as compared to uninfected GSH, indicating cytokine storm and oxidative environment respectively in SARS-CoV2 infection (**Figure S2C)**. Immune activation and infiltration at the target site primarily dependent on chemokines and chemokine receptors. Therefore, we tested the expression of both chemokines and their receptors in SasCov2-infected GSH. We found elevated expression of C-C chemokine receptor (CCR5) and C-C motif chemokine ligand (CCL)-5, at 2 dpi, which regulates T cell function and chemotaxis. CCL-22, which is essential for Treg-DCs cross talk, was found to be elevated at 7 dpi as compared to uninfected GSH. The expression of CXCL9 and CXCL-10, which are crucial for effector T cell trafficking and activation, and have been described as one of the biomarkers for CVC, was upregulated at 2 dpi as compared to uninfected GSH (**Figure 3F)** (Altara et al., 2016; Hueso et al., 2018). In addition, mRNA expression of PD-1 and its ligand, PDL-1 were found to be upregulated at the onset of infection, suggesting the induction of checkpoints to suppress acute immune response induced by SARS-CoV2 infection (**Figure 3G**). These data demonstrated that SARS-CoV2 infection in GSH induces an acute immune activation, chemotaxis and expansion of immune cell population at the peak that resembles a pattern of cytokine storm reported in symptomatic Covid19 patients (Afrin et al., 2020; Moore and June, 2020).

**Figure 3.**
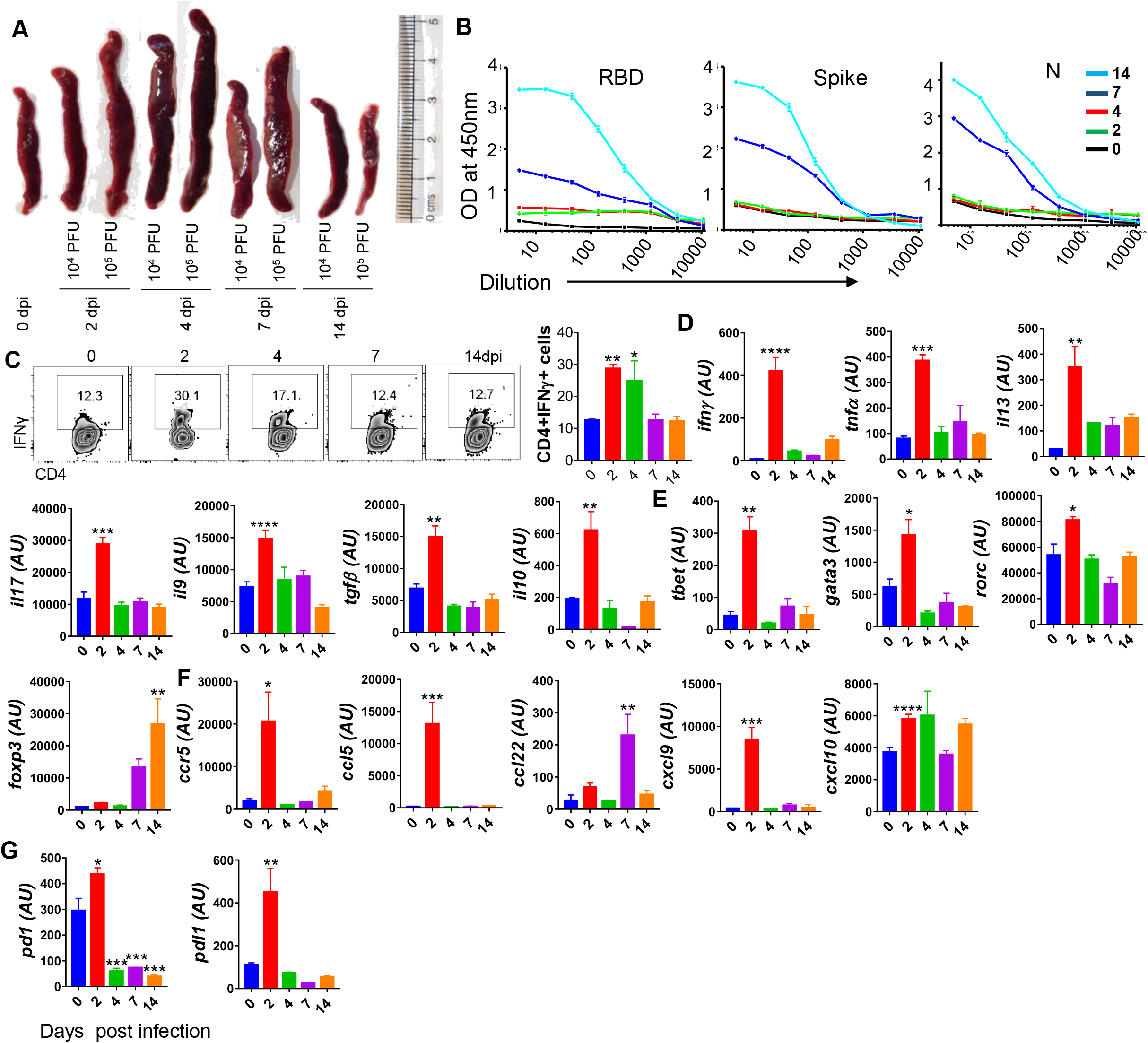
Immunological response against SARS-CoV2 infection in hamsters. (A) Changes in spleen length of SARS-CoV2 infected animals at different dpi as compared to uninfected animals (B) Serum IgG titre against SARS-CoV2 viral proteins (C) FACS for IFNγ secretion showing representative dot plots with % age frequency values and a bar graph showing mean ± SEM. Relative mRNA expression of (D) cytokines, (E) transcription factors, (F) chemokines and (G) checkpoint inhibitors in the spleen of infected vs uninfected hamsters. *P < 0.05, **P < 0.01, ***P < 0.001, ****P < 0.0001 (one-way anova).

### Characterization of cardiovascular pathologies in SARS-CoV2 infected GSH

Myocardial injury was marked in around 25% of hospitalized covid-19 patients that includes thromboembolic diseases and cases of arrhythmia (Giustino et al., 2020; Guo Junyi et al., 2020; Nishiga et al., 2020). The interplay between host-virus interaction through ACE2-RBD and its impact on host renin-angiotensin system (RAS) and immunological response are central to the development of CVC (Giustino et al., 2020). However, due to the lack of a suitable SARS-CoV2 animal model, it has been a limitation for studying cardiovascular-related complications associated with SARS-CoV2 infection. Acute inflammatory response like upregulation CXCL9/10 and oxidative environment allowed to rationalize that SARS-CoV2 infection in GSH may lead to CVC. Indeed, 7 dpi with SARS-CoV2, GSH heart was shown marked ventricular hypertrophy (**Figure S3A).** It was further demonstrated that the ventricular space was found to be reduced characterized by thickening of ventricular walls and interventricular septum at 7 and 14 dpi with SARS-CoV2 as compared to uninfected GSH (**Figure 4A**). The ventricular walls thickening at 7 and 14 dpi with SARS-CoV2 infection in GSH was marked by increased inflammation surrounding the coronary artery and elevated interstitial coronary fibrosis (**Figures 4B, 4C**). Interestingly, increased fibrosis is an established pathophysiological stage in the majority of CVC (Kong et al., 2014; Travers et al., 2016). Further, since CVC are often linked with perturbed serum lipid profile, we reasoned that cardio-vascular pathologies of SARS-CoV2 in GSH may be related to changes in circulating lipid molecules (Bruzzone et al., 2020; D. Wu et al., 2020). Indeed, serum lipid profile showed elevated cholesterol, TGs, HDL, LDL and VLDL levels at 4 dpi (**Figure 4D**). Together, our results provide evidence for CVC arising in hamsters at the later phase of SARS-CoV2 infection.

**Figure 4.**
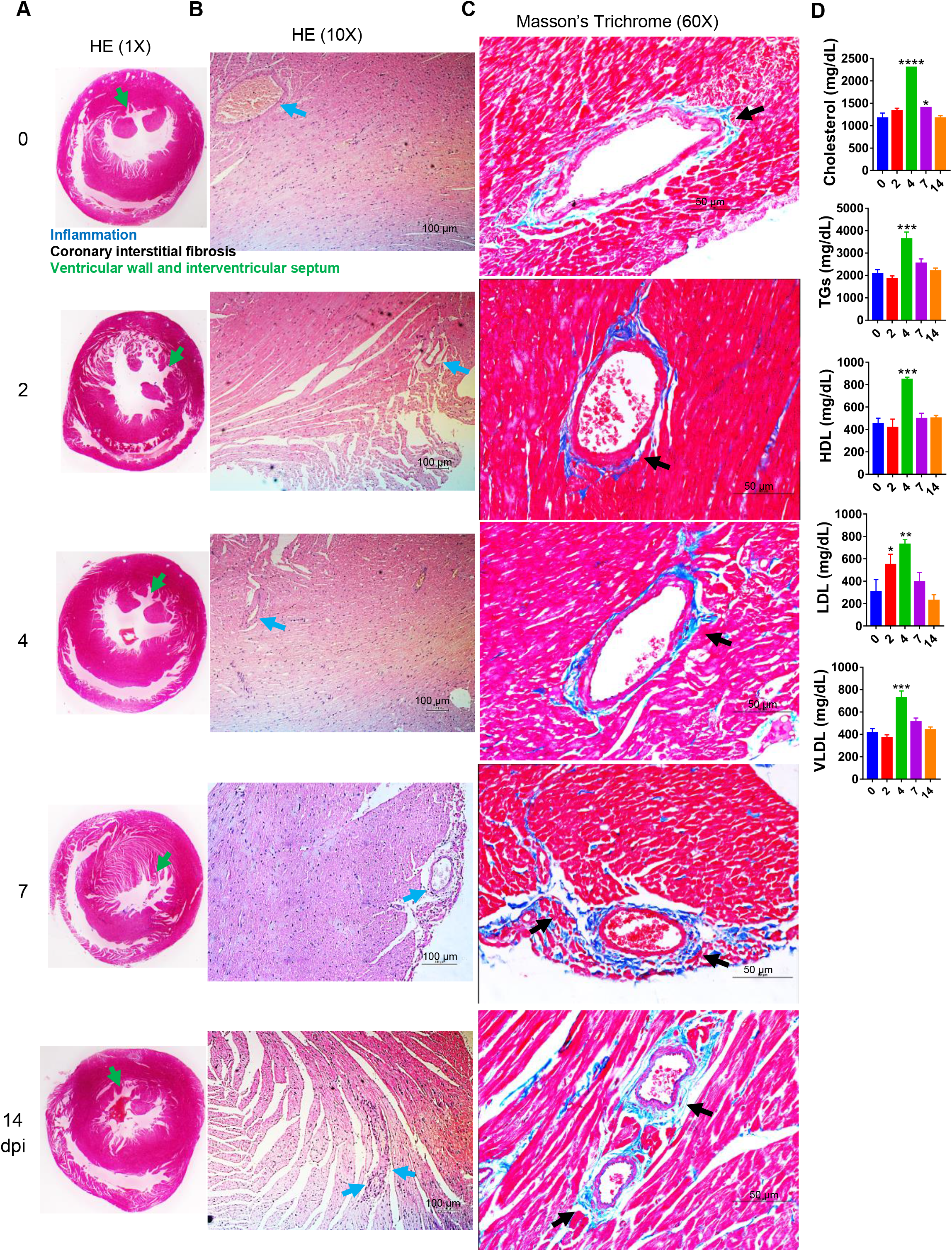
Cardiovascular complications in the heart of SARS-CoV2 infected hamsters. (A & B) heart HE stain captured at 1X and 10X showing ventricular walls and interstitial septum (green arrow) and inflammation around coronary artery (blue arrow) (C) heart MT stains showing interstitial coronary fibrosis (black arrow). (D) serum gross lipid profile. *P < 0.05, **P < 0.01, ***P < 0.001, ****P < 0.0001 (one-way anova).

### Long-chain fatty acid accumulation in SARS-CoV2 infection hamsters may be responsible for CVC

Since our preliminary data revealed CVC and perturbations in lipid profile of SARS-CoV2 infected GSH, we set out to do a detailed characterization of modulation in the abundance of lipid moieties at different time points of infection through ultra-performance liquid chromatography-tandem mass spectrometry (UPLC-MS/MS) to identify the potential lipid biomarkers associated with cardiovascular pathologies. Our LC-MS/MS data identified 831 lipid molecules from the GSH serum samples **(Figure 5A)**. These 831 lipid molecules were from following categories: 11 acylcarnitine (AcCar), 15 cholesterol ester (CE), 50 ceramide (CE), 2 Co-enzyme Q (CoQ), 31 diglycerides (DG), 13 ditetradecanoyl-sn-glycero-3-phosphoethanolamine (DMPE), 10 hexocylceramide (Hex-Cer), 31 lysophosphatidylcholine (LPC), 19 lysophosphatidylethanolamine (LPE), 7 lysophosphatidylinositol (LPI), 105 oxidized (Ox), 107 phosphatidylcholine (PC), 41 phosphatidylethanolamine (PE), 7 phosphatidylglycerol (PG), 38 phosphatidylinositol (PI), 141 ether bond containing lipid (Plasmenyl), 4 phosphatidylserine (PS), 56 sphingomyelin (SM), 5 saturated lipids (SO), 135 triglycerides (TG) (**Figure 5B)**. Interestingly, as compared to uninfected GSH, the lipid profile of 7 dpi with SARS-CoV2 infection showed a distinct PCA distribution plot that corroborated well with the severe cardio-vascular pathology associated with SARS-CoV2 infection at 7 dpi (unpublished data). In order to identify the uniquely modulated lipid molecules on 2, 4, 7 and 14 dpi with SARS-CoV2 in GSH serum samples, we carried out t-test analysis compared to uninfected (0 dpi) control. Volcano plots for 2, 4, 7 and 14 dpi (top to bottom) indicate log2 fold change of lipid molecules which were differentially regulated as compared to the uninfected control (**Figure 5C**). Importantly, 275 lipids were differentially regulated as identified by *t*-test analysis (described in the methods section) of which 65 lipids molecules were up-regulated while 208 were downregulated as identified by log2 fold change by volcano plots (**Figure 5D and Table S1)**. Further, the heatmap for log2 fold change values for differentially regulated lipids and normalized relative abundance showed distinct clusters of lipids at various time points of SARS-CoV2 infection (**Figures 5E, 5F and S3B**). Our lipidomics data show that unsaturated TGs (4-11 double bonds (db)), plasmanly-PE (1-6 db) and plasmanyl-TGs (1-3 db) were uniquely upregulated during infection while saturated TGs (1-3 db), DGs, oxidized lipids, PC, PE and PI were found to be down-regulated. However, both saturated TGs and oxidized lipids, which are implicated in CVC, were downregulated upon infection. Curiously, long-chain fatty acid (LCFAs) TGs were more abundant in infected hamster as compared to uninfected hamsters. Also, medium-chain fatty acid (MCFAs) TGs were more abundant in uninfected hamsters as compared to infected hamsters. that we reasoned as mediators of CVC in GSH as increased LCFAs and decreased MCFAs are associated with increased risks of heart diseases (Labarthe et al., 2008). In addition to CVC, SARS-CoV2 can infect human gastro-intestinal tract cells expressing ACE2 and cause associated pathologies (Lamers et al., 2020). Moreover, severe SARS-CoV2 infection is also known to cause multiple organ failure affecting kidney, liver, brain (Wang et al., 2020). To evaluate extra-pulmonary pathophysiological of infected GSH, we carried out a detailed histological analysis of these major organs. Results show that SARS-CoV2 infection in GSH significantly increases in mucin stain in colons of 2, 4 and 7 dpi, indicative of intestinal inflammation and gastro-intestinal injury (**Figure S4A).** However, there were no gross changes were observed in brain, liver and kidney of SARS-CoV2 infected GSH at 2 dpi as compared to uninfected GSH **(Figure S4B).** Taken together, these data demonstrated a distinct lipidomic profile upon SARS-CoV2 infection.

**Figure 5.**
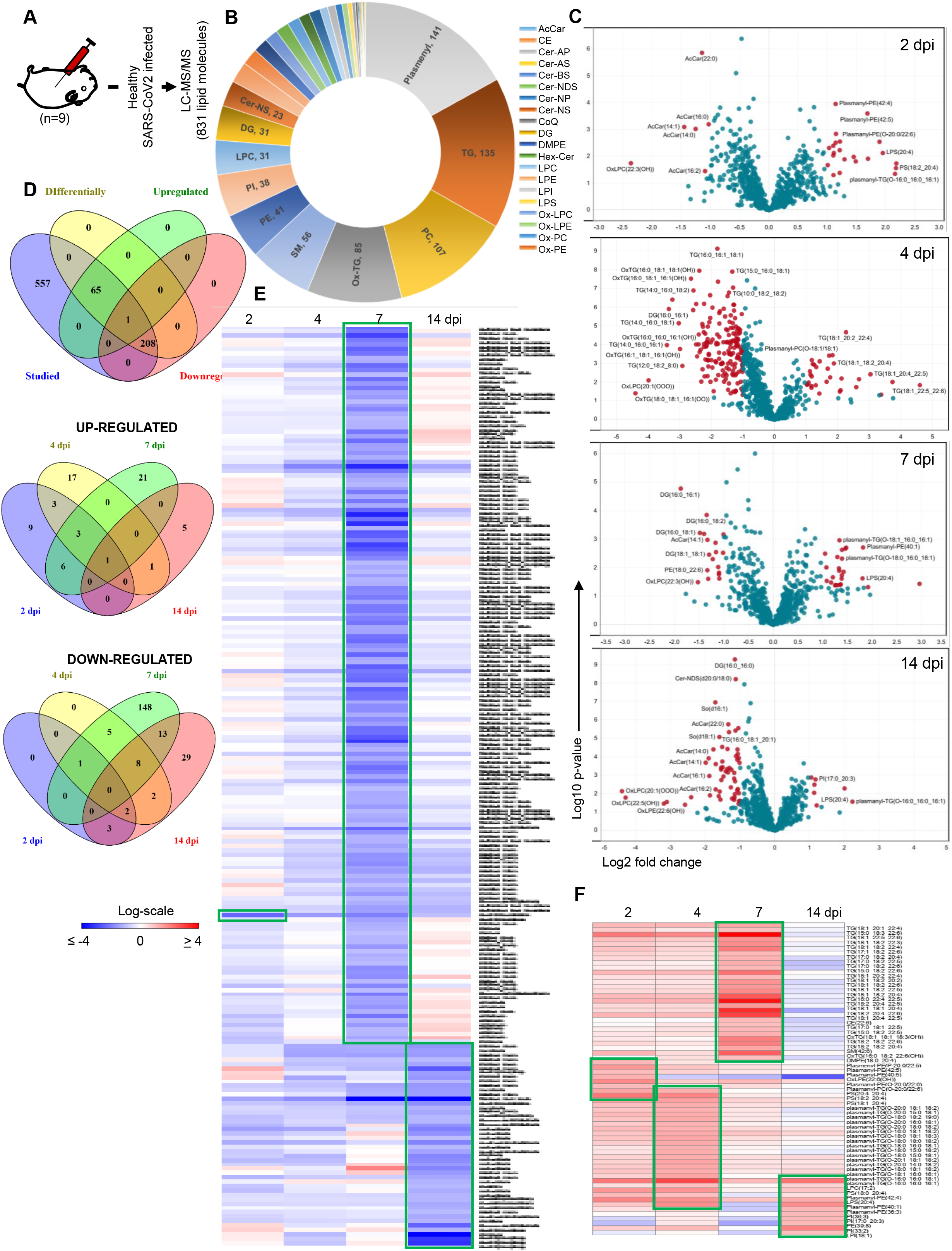
Serum lipid profile of SARS-CoV2 infected hamster. (A) Scheme for lipidomics analysis (B) classification of detected lipid in this study (C) volcano plot for 2, 4, 7 and 14 dpi (D) Venn diagram for differentially regulated lipids (E & F) log2 fold change heatmap.

### Metabolomics changes in SARS-CoV2 infection in hamsters and identification of potential metabolic biomarkers for disease pathophysiology

Emerging literature has suggested that the metabolomic changes are associated with Covid-19 patients (Bruzzone et al., 2020; Grassin-Delyle et al., 2021; Shen et al., 2020). Attempts were made to identify signature metabolites that could help in predicting the severity and progression of Covid-19 disease. In order to find a correlation of metabolic signatures of Covid19 with SARS-CoV2-infected GSH, we carried out metabolomics analysis for 2, 4, 7 and 14 dpi with SARS-CoV2 infected serum samples and compared it with uninfected (0 dpi) control serum samples. A schematic flow for the metabolomics study is shown in (**Figure 6A).***t-test* analysis identified 82 differentially modulated metabolites out of 334, of which 61 metabolites were upregulated while 22 metabolites were downregulated (described in method section) **(Figures 6B, 6C).** The volcano plots indicating log2 fold change in metabolites abundance vs uninfected control was exploited to plot heatmap showing unique clusters (**Figure 6D, E and Table S2**). The relative abundance of individual serum samples has been depicted as a heatmap (**Figure S5B)**. Collectively, riboflavin pathway, arginine and proline metabolism were significantly impacted by these differentially regulated metabolites in SARS-CoV2 infection in GSH (**Figure 6F).** Next in order to understand the correlation of differentially regulated metabolites identified in hamster with the severe and non-severe covid19 patients metabolomics profile we exploited the metabolomics data recently published by *Bo Shen et al 2020* (**Figure 6G)** (Shen et al., 2020). Our correlation study, revealed 21 overlapping metabolites with non-covid 19 metabolites, 37 with non-severe, 37 with severe and 5 with severe vs non-severe metabolites shown as a Venn diagram (**Figures S5C and S5D).** Importantly, 4 metabolites were found to be common between severe covid19 patients and our differentially modulated metabolites viz Citrate, N-acetylneuraminate, allantoin and uracil, while 3 metabolites, Arginine, proline and N-acetylglutamate, were common between severe, non-severe and our differentially regulated metabolites (**Figures 6G, 6H, 6I).** Remarkably, N-acetlyneuraminate and Allantoin were highly upregulated in SARS-CoV2 infection at 2 dpi in our study and corroborated well with severe disease profile in covid19 patients (**Figure 6H)**. Moreover, both N-acetylneuraminate and allantoin have been previously described as CVD biomarkers (Zhang Lei et al., 2018). Together, based on the previously published study we identify and propose biomarkers for disease severity in hamsters and also identify mediators for extra-pulmonary manifestations.

**Figure 6.**
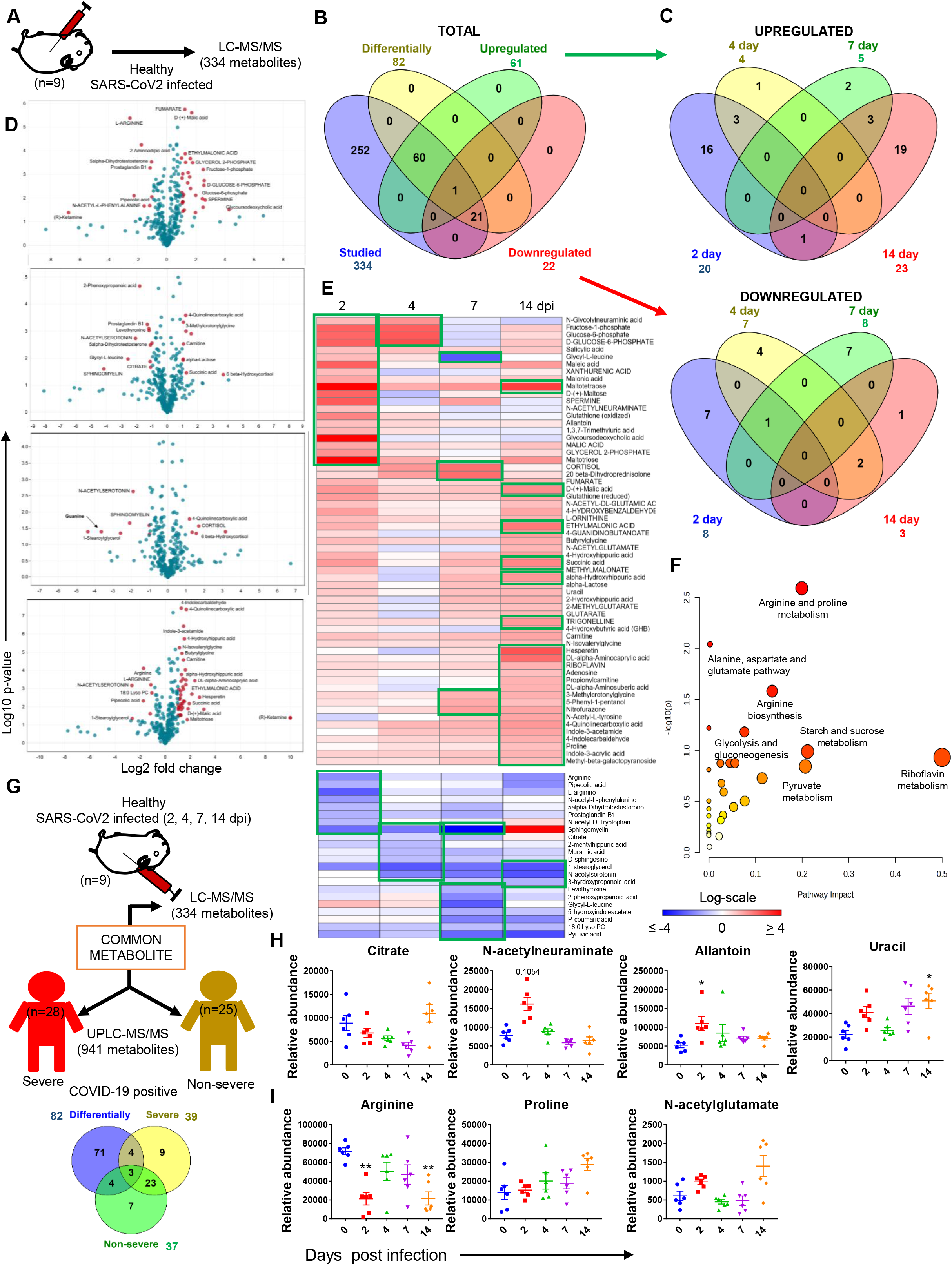
Serum metabolomics profile of SARS-CoV2 infected hamster. (A) Schematic representation for metabolomics. (B & C) Venn diagrams for differentially regulated. (D) volcano plot for 2, 4, 7 and 14 dpi. (E) heatmap for log2 fold change (F) pathways impacted. (G) Scheme and Venn diagram for correlation with Bo Shen et al 2020. (H) dot plot for the relative abundance of metabolites common with the severe profile (I) or severe vs non-severe profile. *P < 0.05 (one-way anova).

## Discussion

Pulmonary, as well as extra-pulmonary clinical and histopathological changes associated with SARS-CoV2 infection in human, had been described by several groups (Chen and Li, 2020; Lamers et al., 2020; Mao et al., 2020; Moore and June, 2020; Neurath, 2020; Nishiga et al., 2020; Verity et al., 2020; Wang et al., 2020, p. 19). GSH as a pre-clinical SARS-CoV2 infection model was recently shown to mimic viral entry and replication naturally similar to human and therefore is often favoured over transgenic mouse model (Chan et al., 2020; Imai et al., 2020; Lee et al., 2020; Sia et al., 2020). The SARS-CoV2 infection in GSH causes lung pathologies, which resembles that of human lung pneumonitis, inflammation and alveolar epithelial injury (Chan et al., 2020; Lee et al., 2020; Sia et al., 2020). This is not surprising as hACE2 and haACE2 receptors had been shown to share major sequence homology (Chan et al., 2020). Such similarities between hACE2 and haACE2 strongly point to interaction with SARS-CoV2 RBD structures with a similar trend of binding affinity. Our *in silico* data on haACE2 extends the previous knowledge providing greater insight into the interaction established between haACE2 and RBD protein. Moreover, the interacting residues and changes in interaction energy is strikingly similar between hamster and human and as a result the furin cleaved S subunit could enter the interacting groove of haNRP-1 and prime it for the interaction with ACE2 in a manner similar to human. Importantly, the host-virus interaction of SARS-CoV2 infection in GSH was found to be similar to human infection for the recently reported SARS-CoV2 mutant. Since, *in silico* data was suggestive of similarity in SARS-CoV2 interaction in hamsters and humans, which corroborates well with intranasal infection and subsequent lung viral load in GSH as reported by previous studies. As increased viral load in the lungs is reflective of lung pathologies and has been shown to induce inflammation and injuries in the clinical cohorts, we reasoned that consistent pathophysiological changes may be occurring in the GSH as well (Leng et al., 2020). Inline, we observed profound pneumonitis, inflammation and alveolar epithelial injury upon histological assessments of the lung. Our data show that lung pathologies long persists after the virus clearance from the lung since we observed a high pathological score on 7 dpi. This corroborates with the clinical cases in which it has been observed that lung pathophysiology is upregulation of the expression of lung injury markers upon SARS-CoV2 infection on 2 and 4 dpi corroborating the lung pathologies.

Viremia is an important pathological characteristic of SARS-CoV2 infection in humans, further SARS-CoV2 is known to shed viral proteins that could be circulating in the serum long after the virus is cleared off (Atkinson and Petersen, 2020). The host responds to these pathogenic factors by producing IgG antibodies for its effective neutralization (Rogers et al., 2020). We found that IgG response to SARS-CoV2 was most profound on 7 and 14 dpi, which corroborates well with recovery of SARS-CoV2 infection in GSH model. In addition to the anti-SARS-CoV2 antibody response, a robust host response against SARS-CoV2 depends on cellular proliferation and immune activation. Consistently, our data demonstrate profound splenomegaly in SARS-CoV2 infected animals on 2 and 4 dpi pointing at active and immune cells stimulation and proliferation. Further, numerous clinical studies have shown that SARS-CoV2 infection is characterized by a cytokine storm, which is responsible for lung pathologies and respiratory distress in severe Covid19 cases. Important mediators of inflammatory response such as IFNγ, IL-6, IL-17A, etc have been shown to be produced in a robust amount as a hallmark of the acute immune response (Moore and June, 2020). Our data show an increased Th1 cytokine, IFNγ at 2 dpi. qPCR studies revealed a similar activation pattern for several inflammatory cytokines and chemokines along with immunoregulatory transcription factors indicating that the immune activation sets in quite early during infection and maybe playing a role in pathological damages associated with lung and other organs during infection.

SARS-CoV2 can affect cardiovascular system which results in inflammation, endothelial activation and microvascular thrombosis (Giustino et al., 2020; Guo Junyi et al., 2020; Nishiga et al., 2020). The CVC is one of the co-morbidity found to be associated with Covid-19 patients. Since there is a lack of knowledge about the CVC arising from SARS-CoV2 infection in GSH, we carried out a detailed histological analysis to monitor the changes occurring in and around cardiomyocytes of the heart muscle. Interestingly, we found profound thickening of the ventricular and interventricular septum with marked depression in ventricular space capacity on 7 and 14 dpi heart samples. These changes were accompanied by the deposition of fibrous mass around the coronary artery and inflammation, which is believed to be one of the major causes of CVC and arrhythmia (Travers et al., 2016). Observation of cardio-vascular complication on 7 and 14 dpi was a remarkable finding, as it hinted strongly at extra-pulmonary damages caused by SARS-CoV2 infection. It also pointed at soluble mediators of the host, which could lead to CVC long after the virus is cleared off from the lungs. One such important mediators of cardio-vascular diseases are serum lipid molecules (Labarthe et al., 2008). We found global changes in the lipid profile associated with infection but quite remarkably 7 and 14 dpi samples produced the largest clusters of altered lipid molecules which were in perfect agreement with heart histology data. Interestingly, SARS-CoV2-infected GSH serum samples were shown an increase in LCFAs accumulation as compared to MCFAs, which have been described as biomarkers of CVC and heart failure.

In addition to the lack of data on the extra-pulmonary changes associated with SARS-CoV2 infection GSH, no studies have been shown to evaluate the metabolomics changes occurring in the GSH infected with SARS-CoV2. We observed striking similarities between the metabolomics profile of GSH compared with the clinical database (Shen et al., 2020). When compared to the metabolomic profile of severe and non-severe Covid19 patients, 7 metabolites were found to be common in GSH. Both allantoin and N-acetylneuraminate increased abundance have been shown to be biomarkers for increased heart risk (Maciel et al., 2016; Zhang Lei et al., 2018). Together, our data provide the first insight into the key metabolites which could be used as the biomarkers for the efficient and rapid evaluation of SARS-CoV2 disease severity.

As of now hamster model for SARS-CoV2 has been described as a suitable model to study lung pathologies and associated clinical parameters (Chan et al., 2020; Imai et al., 2020; Kaptein et al., 2020; Kreye et al., 2020; Lee et al., 2020; Osterrieder et al., 2020; Sia et al., 2020; Tostanoski et al., 2020). However, a robust animal model that mimics the extrapulmonary pathophysiological changes upon SARS-CoV2 infection and resembles long Covid symptoms has not been evaluated. The current study provides a detailed insight into the cardio-vascular and immunopathological changes associated with SARS-CoV2 progression in GSH. Our data further demonstrate distinct lipidomic and metabolomics changes in SARS-CoV2 infected GSH. Together, our study provide the first proof-of-concept (PoC) for a robust preclinical model to evaluate both early and late consequences of SARS-CoV2 infection.

## Acknowledgments

Financial support to AA laboratory from THSTI core and Translational Research Program (TRP) and Department of Biotechnology (DBT) and DST-SERB. We acknowledge IDRF (THSTI) for the support at ABSL3 facility. Luvas (University of Hisar) and CDRI for providing the hamster for the study. Small animal facility and Immunology Core for providing support in experimentation. ILBS for support in histological analysis and assessment. RCB microscopy facility for microscopic examination of histology slide. We thank Dr. Gagandeep Kang, CMC Vellore for providing support for this project. The following reagent was deposited by the Centers for Disease Control and Prevention and obtained through BEI Resources, NIAID, NIH: SARS Related Coronavirus 2, Isolate USA-WA1/2020, NR-52281. AKY is supported by DBT-Big Data Initiative grant (BT/PR16456/BID/7/624/2016) and the Translational Research Program (TRP) at THSTI funded by DBT.

## Author Contributions

Conceived, designed and supervised the study: AA; Designed and performed the experiments: AA, ZAR, RD, SS; ABSL3 procedures: ZAR, RD, SS, AKP; *in silico* study: SA; qPCR primer designing and analysis: ZAR; FACS: ZAR; Histology and analysis: ZAR, MRT; Metabolomics: YK, ZAR, SKG; Lipidomics: YK, ZAR, SKG; ELISA: TS; Viral load: GPM; Virus stock and TCID50: SSamal; Omnics analysis: AKY, SAgarwal, ZAR, RD, SS; Contributed reagents/materials/analysis tools: AA; Wrote the manuscript: ZAR, AA.

## Star Methods

### Animal Ethics and biosafety statement

6-8 weeks old female golden Syrian hamsters were acclimatized in biosafety level-2 (BSL-2) for one week and then infected in Animal BSL3 (ABSL-3) institutional facility. The animals were maintained under 12 h light and dark cycle and fed standard pellet diet and water ad libitum. All the experimental protocols involving the handling of virus culture and animal infection were approved by RCGM, institutional biosafety and IAEC animal ethics committee.

### Virus preparation and determination of viral titers

SARS-Related Coronavirus 2, Isolate USA-WA1/2020 virus was used as challenge strain, which was grown and titrated in Vero E6 cell line grown in Dulbecco’s Modified Eagle Medium (DMEM) complete media containing 4.5 g/L D-glucose, 100,000 U/L Penicillin-Streptomycin, 100 mg/L sodium pyruvate, 25mM HEPES and 2% FBS. The virus stocks were plaque purified and amplified at THSTI Infectious Disease Research Facility (Biosafety level 3 facility) as described previously (Harcourt et al., 2020; Mendoza et al., 2020).

### SARS-CoV2 infection in golden Syrian hamster

Infection in GSH was carried out as previously described (Chan et al., 2020; Sia et al., 2020). Briefly anesthetized through ketamine (150mg/kg) and xylazine (10mg/kg) intraperitoneal injection and thereafter infection was established intranasally with 10^4^ PFU (100μl) or 10^5^PFU (100μl) of live SARS-CoV2 or with DMEM mock control inside ABSL3 facility.

### Clinical parameters of SARS-CoV2 infection

All infected animals were housed for 14 days and their body mass was monitored as previously described (Chan et al., 2020; Sia et al., 2020). 9 animals from each group were sacrificed on 2, 4, 7 and 14 dpi and their blood serum along with body organs such as lungs, intestine, liver, spleen and kidney were collected. Serum samples were stored at −80 °C until further use. Lung and spleen samples of infected and uninfected animals were compared for any gross morphological changes. Lungs samples were homogenized in 2 ml DMEM media and used for viral load and TCID50 determination. A section of the lung, intestine along with other organs were fixed in 10% formalin solution and used for histological studies. Spleen was strained through 40 μm cell strainer with the help of a syringe plunger and used for qPCR and immunophenotyping studies.

### Viral load

Homogenized lung samples of 2, 4, 7, 14 dpi along with uninfected controls were centrifuged for 10 min at 4°C and their supernatant was collected. 100 μl of supernatant from each sample was then mixed with 900 μl of Trizol reagent (Invitrogen) and RNA isolation was carried out as per the manufacturer’s protocol. Copy number estimation of SARS-CoV-2 RNA has been described previously (Anantharaj et al., 2020). Briefly, 200 ng of RNA was used as a template for reverse transcription-polymerase chain reaction (RT-PCR). The CDC-approved commercial kit was used for of SARS-CoV-2 N gene: 5′-GACCCCAAAATCAGCGAAAT-3′ (Forward), 5′-TCTGGTTACTGCCAGTTGAATCTG-3′ (Reverse) and 5′-FAM-ACCCCGCATTACGTTTGGTGGACC-BHQ1-3′ (Probe) detection and sub-genomic RNA copy numbers were estimated by ΔΔCt method. Hypoxanthine-guanine phosphoribosyltransferase (HGPRT) gene was used as an endogenous control for normalization through quantitative RT-PCR. The region of N gene of SARS-CoV-2 starting from 28287 – 29230 was cloned into pGEM®-T-Easy vector (Promega). This clone was linearized using SacII enzyme and *in vitro* transcribed using the SP6 RNA polymerase (Promega). The transcript was purified and used as a template for generating a standard curve to estimate the copy number of SARS-CoV-2 N RNA (Anantharaj et al., 2020).

### TCID50

For TCID50 determination, 50 μl of homogenized lung supernatant samples were incubated with confluent Vero-E6 cells in 96-well plates as described previously (Chan et al., 2020). Briefly, serial dilutions of 10-folds from each sample were added to the wells containing Vero-E6 cells monolayer in DMEM media in quadruplicate. After 4 days of incubation, TCID50 was determined through Reed and Münch endpoint method with one TCID50 equivalent to the amount of virus required to cause cytopathic effect in 50% of inoculated wells. TCID50 values were expressed as TCID50/ gram of lung mass. For the determination of virus titers, tissue samples (lungs) were homogenized to a final 10% (w/v) suspension in DMEM medium with gentamicin (Invitrogen, USA). The tissue samples from 2 to 14 dpi were used to infect Vero E6 cell monolayers in 48-plates as described previously. Virus titers were expressed as TCID_50_ per gram of tissue

### ELISA

The binding-antibody response to SARS CoV-2 post-infection was measured using ELISA based platform as described earlier (Shrivastava et al., 2018). The antibody response was measured against Spike, RBD and N protein post virus challenge at different time points. Soluble spike prefusion trimeric protein (S2P) (Wrapp et al., 2020), in house RBD (Ha et al., 2020) and N proteins were coated on Maxisorp plates (Nunc) at different protein concentrations (Spike S2P trimer; 1 μg/ml, RBD; 2μg/ml and N protein at 0.5ug/ml) in 1X carbonate/bicarbonate buffer, pH 9.6 overnight at 4 °C. The following day, the plates were blocked using 250 μl of PBS containing 5 % skimmed milk (blocking buffer). Three-fold serially diluted sera (with 1:5 as starting dilution) in dilution buffer (1:5 times dilution of blocking buffer) were added to wells of the plates. The plates were incubated at room temperature (RT) for 1 h and then washed three times with washing buffer (PBS + 0.05 % tween 20), the ELISA plates with N protein coating was washed additionally once with high salt PBST (Phosphate buffer with 500 mM NaCl and 0.05 % Tween 20) and incubated with biotinylated anti-hamster IgG antibody (Sigma) for another 1 h and washed subsequently with the washing buffer and incubated further with Avidin-HRP (Sigma) for 45 min at RT. Post incubation plates were washed four times and 100 μl of TMB substrate (Thermo Fisher Scientific) was added to the washed wells. The reaction was stopped by adding 100 μl of 1 N H_2_SO_4_ and the plates were read at 450 nm on a 96-well microtiter plate reader. Sera end point titers were calculated as the reciprocal of serum dilution giving OD 450 nm readings higher than lowest dilution of the placebo or control arm + two times standard deviations

### *In silico* molecular modeling and protein-protein docking

In the absence of crystal structure of hamster ACE2 structures, the crystal structure of human ACE2 and RBD, pdb-ids 6m17, 6VW1 were considered as templates to generate the robust model of hamster through homology modeling (Srivastava et al., 2018). The robustness of each model was evaluated through ERRAT, PROSA and Ramachandran plot (allowed and disallowed %), followed by structure optimization through energy minimization to allow conformational relaxation of the protein structure (Ha et al., 2020). The same was carried out for neurolipin-1 protein of hamster as human neurolipin crystal structure (pdb-id 6JJQ) is reported. The most stable proteins were picked to perform the protein-protein (for RBD-ACE2) and protein-peptide (NRP-1 and CendR peptide of spike protein) docking studies through different algorithms. Rigid docking through PyDock allows some steric clashes while flexible docking by SwarmDock was done on relaxed structures of proteins to generate candidate solutions with at least one near-native structure (Mattapally et al., 2018). Clustering of docked poses was conducted to generate the most likely complex of ACE2-RBD and NRP1-Cend peptide based on the number of conformers and lowest binding energy (from −50 to −40 kcal/mol). Post-docking analysis followed by energy minimization of complexes showed that both the protein structures displayed an initial structural rearrangement that was followed by convergence indicating structural fluctuation at interface site especially at residual level i.e conformational changes and chi angle variation. The overall structural fluctuation of hACE2 (human ACE2) was observed to be less than that of haACE2 (hamster ACE2). This is expected as the protein structure (template) of the former has high resolution and covers more space, while the latter is the model. An inventory of structural and energetic features of the complexes was obtained by analysing the hydrogen bonds (cut-off of 3.5Å for the donor-acceptor distance and 150° for the donor-hydrogen-acceptor angle) and hydrophobic contacts (non-polar atoms separated by a distance of at-most 4.0 Å). π-π interactions were considered to be formed when the short inter-atomic carbon-carbon distance (SICD) was smaller than 4.8 Å (Kanwal et al., 2016). In an effort to dissect these interactions from the docking simulations, total interaction energy between ACE2 and RBD was calculated, within the framework of Amber-force field description.

### qPCR

RNA was isolated from lung homogenate and spleen samples using Trizol-choloroform as previously described (Rizvi et al., 2018). Thereafter, RNA was quantitated by NanoDrop and 1 μg of total RNA was then reverse-transcribed to cDNA using the iScript cDNA synthesis kit (Biorad; #1708891) (Roche). Diluted cDNAs (1:5) was used for qPCR by using KAPA SYBR® FAST qPCR Master Mix (5X) Universal Kit (KK4600) on Fast 7500 Dx real-time PCR system (Applied Biosystems) and the results were analyzed with SDS2.1 software. The relative expression of each gene was expressed as fold change and was calculated by subtracting the cycling threshold (Ct) value of hypoxantine-guanine phosphoribosyltransferase (HGPRT-endogenous control gene) from the Ct value of target gene (ΔCT). Fold change was then calculated according to the previously described formula POWER(2,-ΔCT)*10,000 (Malik et al., 2017). The list of the primers is provided as follows.

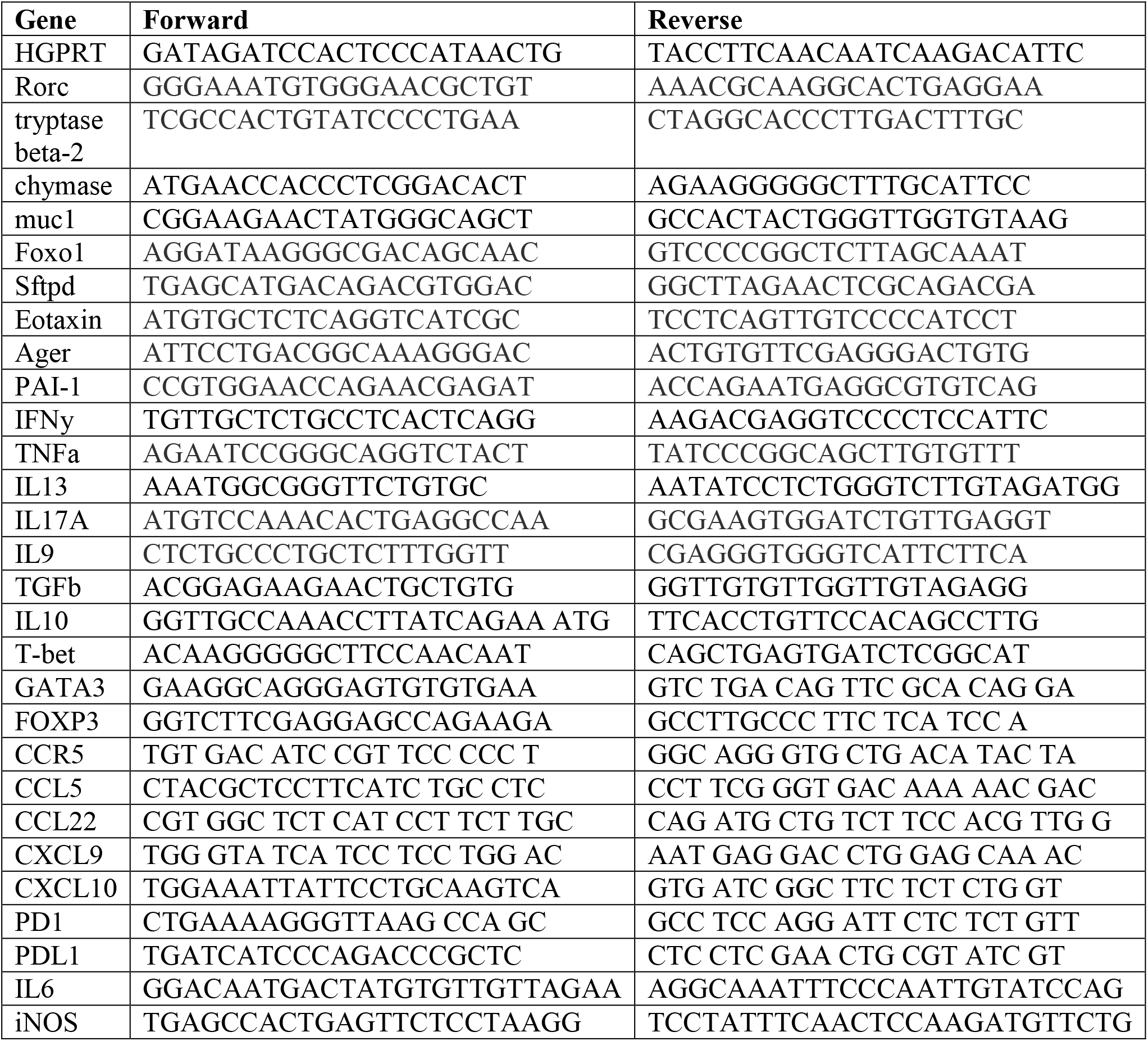

### Histology

Excised tissues of animal organs were fixed in 10% formaline solution and processed for paraffin embedding. The paraffin blocks were cut into 3-μm-thick sections and then mounted on silane-coated glass slides. One section from each organ sample was stained with hematoxylin and eosin. Lung, heart and colon samples were stained with taulidine blue, masson’s trichrome and mucicarmine stains respectively. Each stained section was analyzed and captured at 10X and 60X magnification. Heart images were also captured at 1X magnification. Blind assessment and scoring of each section for each samples were performed by a professional histologist.

### Immunophenotyping of splenocytes

0.5 million RBC lysed splenocytes were used for intracellular cytokine staining by re-stimulation with PMA (phorbol 12-myristate13-acetate; 50 ng/ml; Sigma-Aldrich), ionomycin (1.0 μg/ml; Sigma-Aldrich) and monensin (#554724; GolgiStop, BD Biosciences) for 6 h or with RBD protein (50 μg) for 72 h. Cell surface staining with anti-mouse CD4 (GK1.5) PerCp (Biolegend) was carried out for 20 min in dark at RT. Intracellular anti-mouse IFN-γ (XMG1.2) (BioLegend) staining was then carried out after fixing the cells in Cytofix solution and permeabilization with 1X Perm/Wash Buffer using kit (BD Biosciences; #554714) for 20 min in dark at RT. Cells were then washed and acquired on FACS Canto II and were analyzed with FlowJo software (Tree star) as previously described (Malik et al., 2017).

### Lipid extraction and lipidomics

Lipid extraction from serum sample was performed as previously described method with some modifications (Schwaiger et al., 2018). 0.3 ml methanol was added to 100 μl of serum samples and mixed for 30 sec followed by the addition of 1.25 ml methyl-tert-butyl ether (MTBE). The mixture was incubated at RT for 1 h on a shaker followed by the addition of 0.3 ml of MS grade water was for the phase separation. 10 min after incubation at RT, samples were spin down at 400 rpm at 10 °C for 5 min. Organic upper phase was collected and dried in a speed vac and stored at −80 °C till further use. Moreover, lipids extracted were dissolved in 100 μl of 65:30:5 (acetonitrile: 2-propanol: water v/v/v). An acquity HSS T3 (2.1 mm X 100 mm X 1.8 Um, WATERS) was then utilized for carrying out lipid separation by exploiting ultra-performance liquid chromatography (UPLC). Solvent A and B were water/ acetonitrile (2:3 v/v) and 2-propanol/acetonitrile (9:1, v/v) respectively, at 0.3ml/min flow rate and 40 °C column temperature. 18 min total run time was utilized with the following gradient setup, 0 to 12 min solvent B ramped from 30 to 97% and hold for another 3 min. From 15.2 to 18 min, B was at 30%. Acquisition of data was carried out on a high-resolution mass spectrometer, orbitrap fusion (Thermo Scientific) equipped with a heated electrospray ionization source. Auxiliary gas and ESI sheath gas were 20 and 60 respectively. The negative and positive spray voltage was 3000 volts. For a full MS run, 120k resolution was used with automatic gain control (AGC) targeted of 200000 and mass ranges between 250-1200. The resolution was kept at 30K with AGC target 50000for MSMS. Collision energy for fragmentation used was 27+/−3.

### Lipid data analysis

Lipidmatch flow was utilized with default settings for peak picking (using mzmine), lipid annotation, blank filtration and combining positive and negative data (Koelmel et al., 2017). Thereafter, data analysis was carried out as follows: the data was normalized by sum, perato scaled, and log-transformed for analysis in metaboanalyst.

### Metabolomics analysis LC-MS/MS reverse phase and HILIC

Metabolites were extracted from 100 μl of serum samples by using 100% methanol. Thereafter, mixture was vortexed for 1 min and kept on ice for protein precipitation. After centrifugation (10000 rpm for 10 min at 4 °C) supernatant was collected and distributed in two tubes for polar and nonpolar metabolite analysis. Collected supernatants were then dried using a speed vacuum at for 20 to 25 min at RT and stored at −80 °C till further analysis. For the reverse phase metabolites were dissolved in 15% methanol in water (v/v) and for polar metabolite analysis samples were dissolved in 50% acetonitrile in water (v/v).

### Measurement of metabolites

Orbitrap Fusion mass spectrometer (Thermo Scientific) coupled with heated electrospray ion source was used for data acquisition. Data acquisition methods have been followed as per published protocols (Kumar et al., 2020; Naz et al., 2017) with minor modifications. Briefly for MS1 mode, mass resolution was kept at 120,000 and for MS2 acquisition, mass resolution was 30,000. Mass range of data acquisition was 60–900 da. Extracted metabolites were separated on UPLC ultimate 3,000. Data were acquired on reverse phase and HILIC column and positive and negative ionization mode both. Reverse phase column was HSS T3 and HILIC column was XBridge BEH Amide (Waters Corporation). For polar compound separation, solvent A was 20 mM ammonium acetate in the water of PH 9.0 and mobile phase B was 100% acetonitrile. The elution gradient starts from 85% B to 10% B over 14 min with flow rate of 0.35 ml/min. For reverse phase, Solvent A was water and B was methanol with 0.1% formic acids added in both. The elution gradient starts with 1% B to 95% B over 10 min with flow rate 0.3 ml/min. sample injection volume was 5ul. Pool quality control (QC) sample was run after every five samples to monitor signal variation and drift in mass error. Data processing. All LC/MS acquired data has been processed using the Progenesis QI for metabolomics (Water Corporation) software using the default setting. The untargeted workflow of Progenesis QI was used to perform retention time alignment, feature detection, deconvolution, and elemental composition prediction. Metascore plug of Progenesis QI has been used for the in-house library with accurate mass, fragmentation pattern and retention time for database search. We have also used an online available spectral library for further confirmation of identification. Cut-off for retention time match was 0.5 min and spectral similarity was more than 30% fragmentation match in Progenesis QI. Peaks that had a coefficient of variation (CV) less than 30% in pool QC samples were kept for the further analysis of data. Additionally, manual verification of each detected feature has been done for the selection of right peaks.

### Lipidomics and metabolomics data analysis

For both metabolomics and lipidomics data, the data was cleaned up to remove zero intensity values. The average calculations were done with the remaining available intensity values. The fold change was calculated as the ratio of average intensity values of cases divided by the average intensity of controls as-

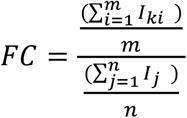

Where,

*FC = fold change*

*m = number of available non-zero intensity values for case*

*n = number of available non-zero intensity values for control*

*I_ki_ = intensity of i^th^ case sample for condition k, where k ∈ {2D, 4D, 7D, 14D post infection}*

*I_j_ = intensity of j^th^ control sample*

A two-tailed t-test considering similar means of case and control population was used to calculate the p-values. The FC was transformed to log_2_FC and a threshold was set to ±2 FC (±1 at log_2_FC). The significance threshold for differentially expression was set using simultaneous threshold filters of the log_2_FC of ±1 as well as a p-value threshold of ≤0.05. This was followed for both the metabolomics and the lipidomics data. The significantly expressed analytes (metabolites and the lipids) are also represented on the volcano plots. For the volcano plots, the p-values were transformed into –log_10_ P-values for plotting. The significant values as described above were highlighted and labeled on the graph using Tableau (version 2020.3). **Volcano plot** -log_10_P value vs log_2_ fold change was plotted on the Volcano plot. Tableau (version 2020.3) was used to highlight and annotate some of the significant hits(metabolites/lipids). The points above or below ±1 log_2_ fold change (i.e. ±2 fold change) and 1.33 on the –log_10_Pvalue scale (i.e. ≤ 0.05 p-value) were selected for annotation and were colored red for visual distinction. For the heat map construction, we used log_2_ fold changes for both the metabolites as well as lipids heat maps. Gene-E software (https://software.broadinstitute.org/GENE-E/) was used to create the heat maps. For the analyte (metabolite/lipid) with a zero values in control but a positive intensity reading value in cases, we considered them as overexpressed vs control in that particular condition, and used a maximum log_2_ FC imputed manually. The value chosen for such cases was the maximum value from the data. For setting the color scale, we used blue color to represent all the values below −4 log_2_ FC (1/16 FC) and all the values above +4 log_2_ FC were colored red (+16 FC). **For whole data heatmap** Clustvis was used to generate the heat map for the complete dataset for both metabolites as well as lipids. **PCA plots** raw values of the metabolites and lipids were taken for all 9 samples. The web-based tool ClustVis was used to create the sample PCA plots to check the clustering of biological samples (ClustVis: a web tool for visualizing clustering of multivariate data using Principal Component Analysis and heatmap) (Metsalu and Vilo, 2015). **Comparison against the human data** The data from article (Shen et al., 2020) was downloaded and manually compared with the hamster metabolites data obtained from our study. The metabolites common to each category were then compared against each other to identify metabolites unique to each category as classified by the authors.

### Statistical analysis

All the results were analyzed and plotted by using Graph pad prism 7.0 software. Body mass, lung mass, gene expression, FACS, ELISA, TCID_50_ and qPCR studies were compared and analyzed by using one-way ANOVA or student t-test. Metabolomics and lipidomics analysis was carried out with n = 9 serum samples per group. P-value of less than 0.05 and was considered as statistically significant.

## Supplementary tables and legends

**Table S1. Differentially regulated lipids molecules at different time points.**

**Table S2. Differentially regulated metabolites at different time points.**

## Supplementary figures

**Figure S1.***In silico* **analysis.** (A) Scheme for h/ha ACE2-RBD interaction (B) catalytic triad for hamster TMPRSS2.

**Figure S2. Immunological response.** (A) antibody titre for different dose of infection (B) FACS dot plot and bar graph showing mean ± SEM % frequency of CD4+ cells. (C) bar graph showing mean value ± SEM of mRNA expression of IL6 and iNOS genes in the splenocytes samples. ****P < 0.0001 (one-way anova).

**Figure S3. Morphological changes in heart and lipid relative abundance heatmap.** (A) Heart hypertrophy on 7 dpi (B) heatmap for normalized relative abundance.

**Figure S4. Extra-pulmonary changes** (A) H & E stain (B) mucicarmine stain for the presence of mucin (blue arrow) of colon on 2, 4, 7 and 14 dpi (C) H & E stain for brain, liver and kidenty on 2 dpi. ***P < 0.001, ****P < 0.0001 (one-way anova).

**Figure S5. Metabolomics.** (A) PCA plot for metabolomics (B) heatmap showing normalized relative abundance (C & D) Venn diagram showing correlation of metabolites with Bo Shen et. al. metabolomics profile in covid19 patients.

---

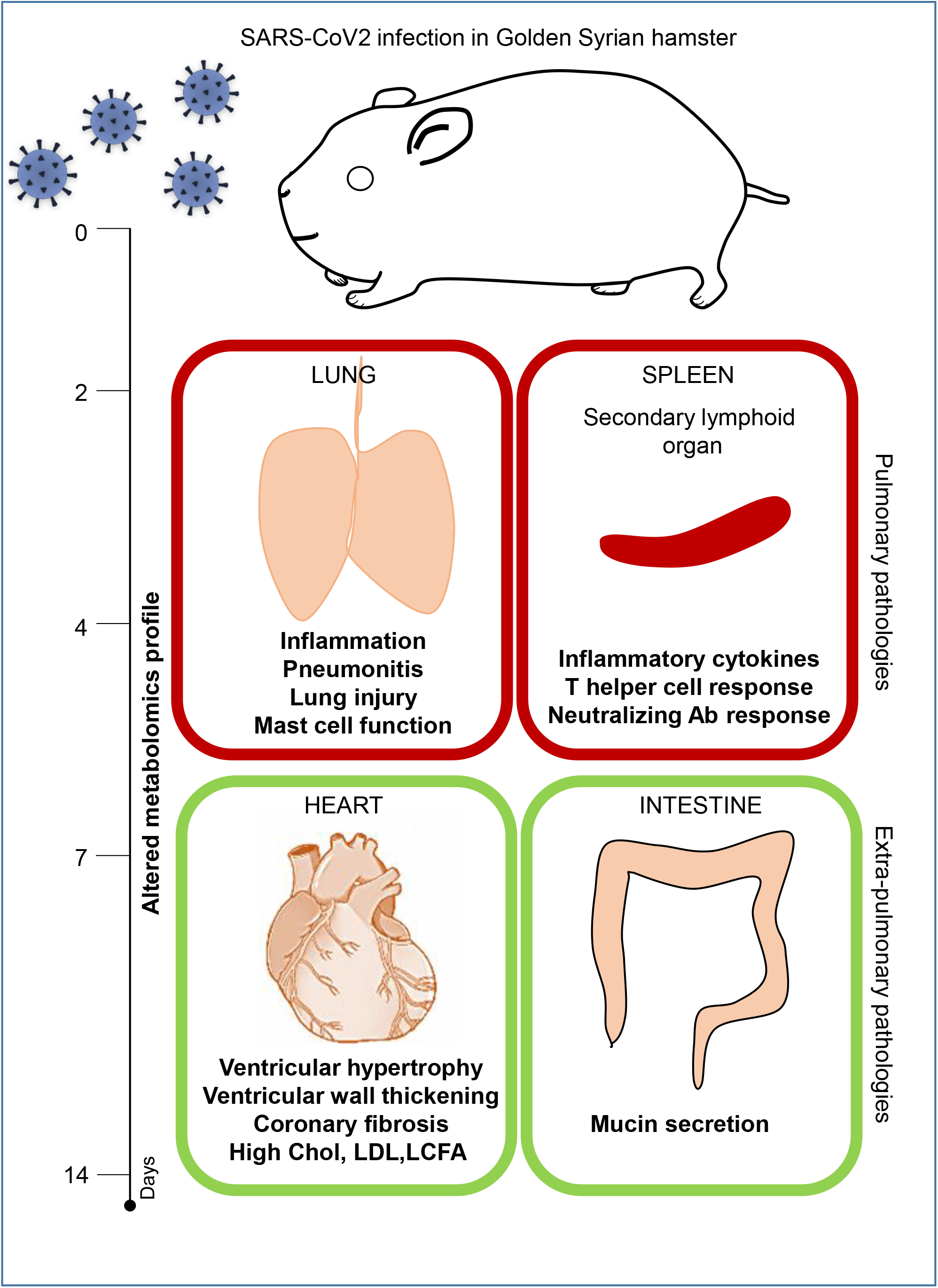

