## Supplementary figures and images for "Immunological and cardio-vascular pathologies associated with SARS-CoV-2 infection in golden syrian hamster"

### Supplementary figures

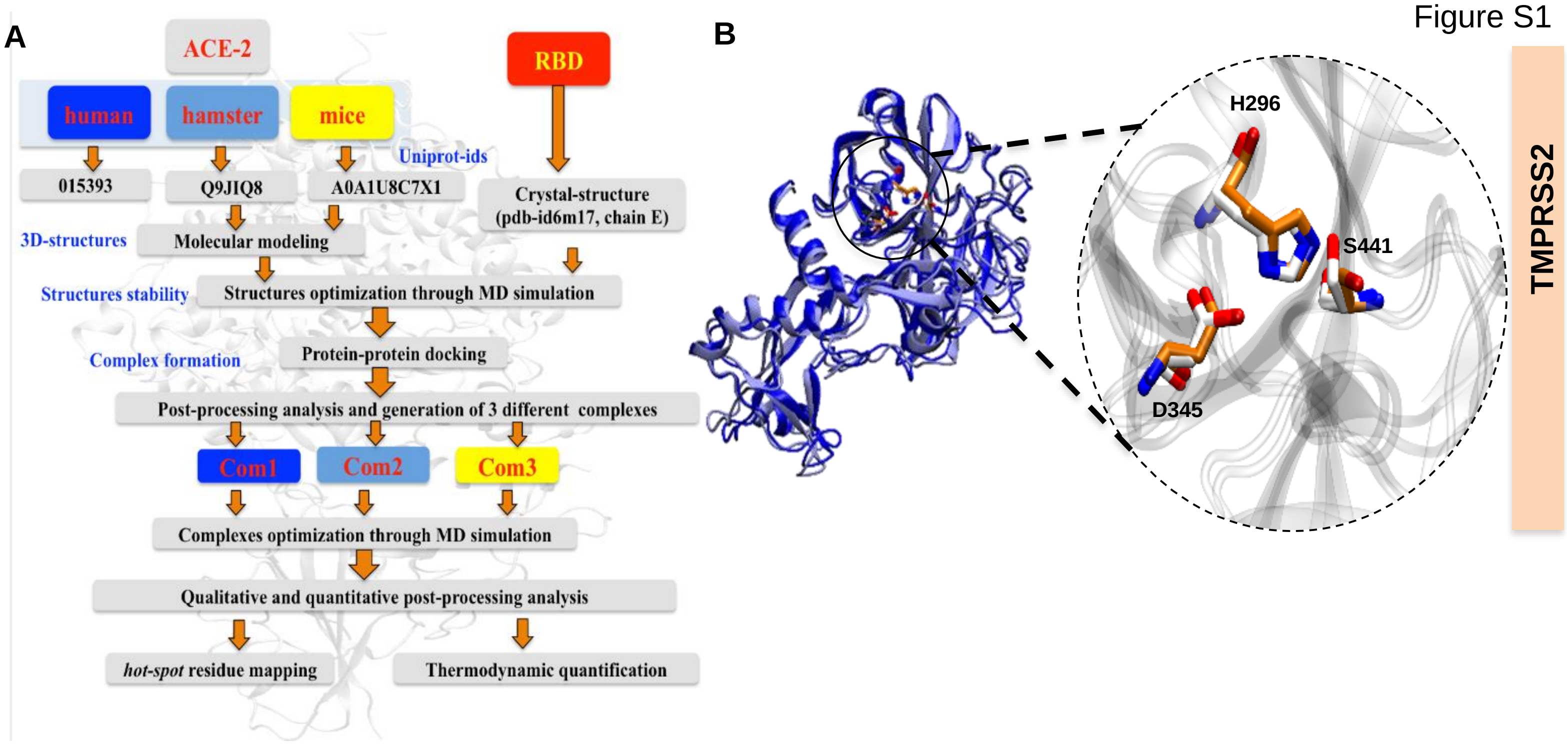

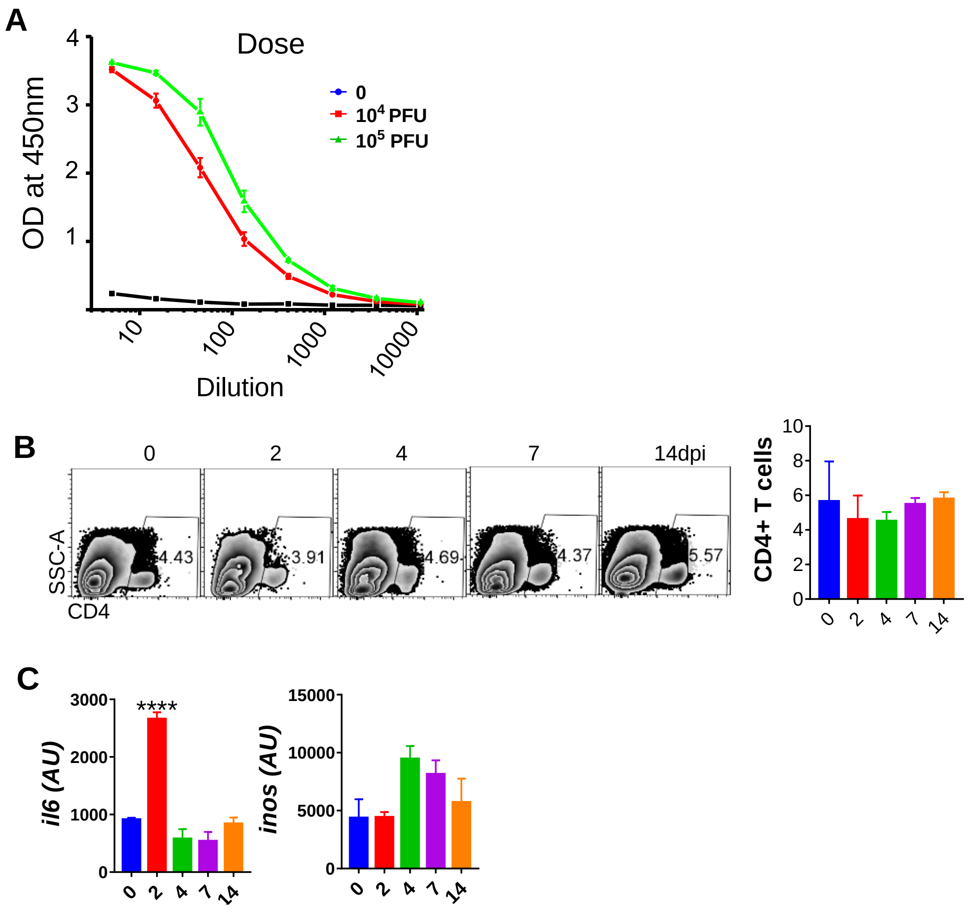

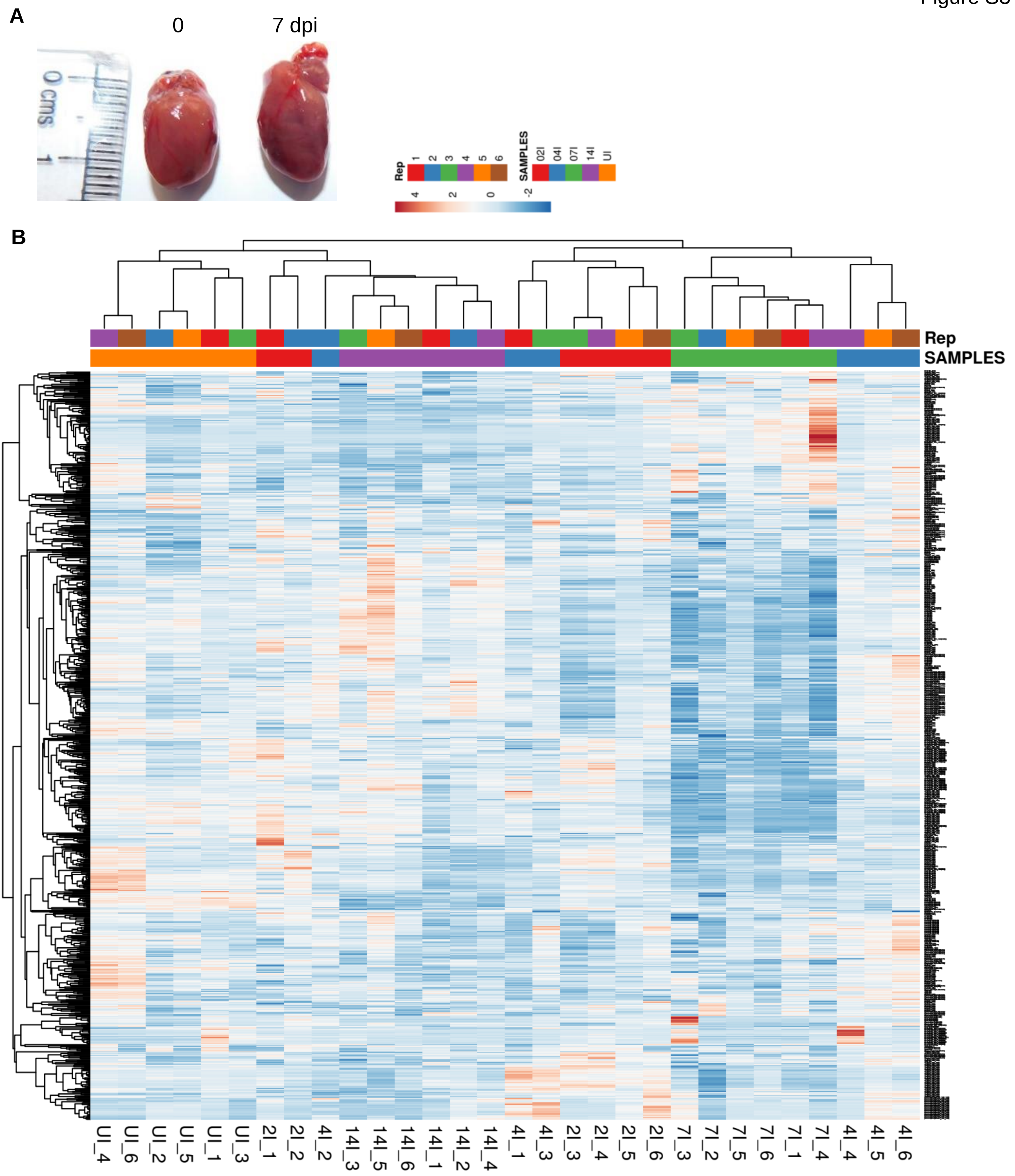

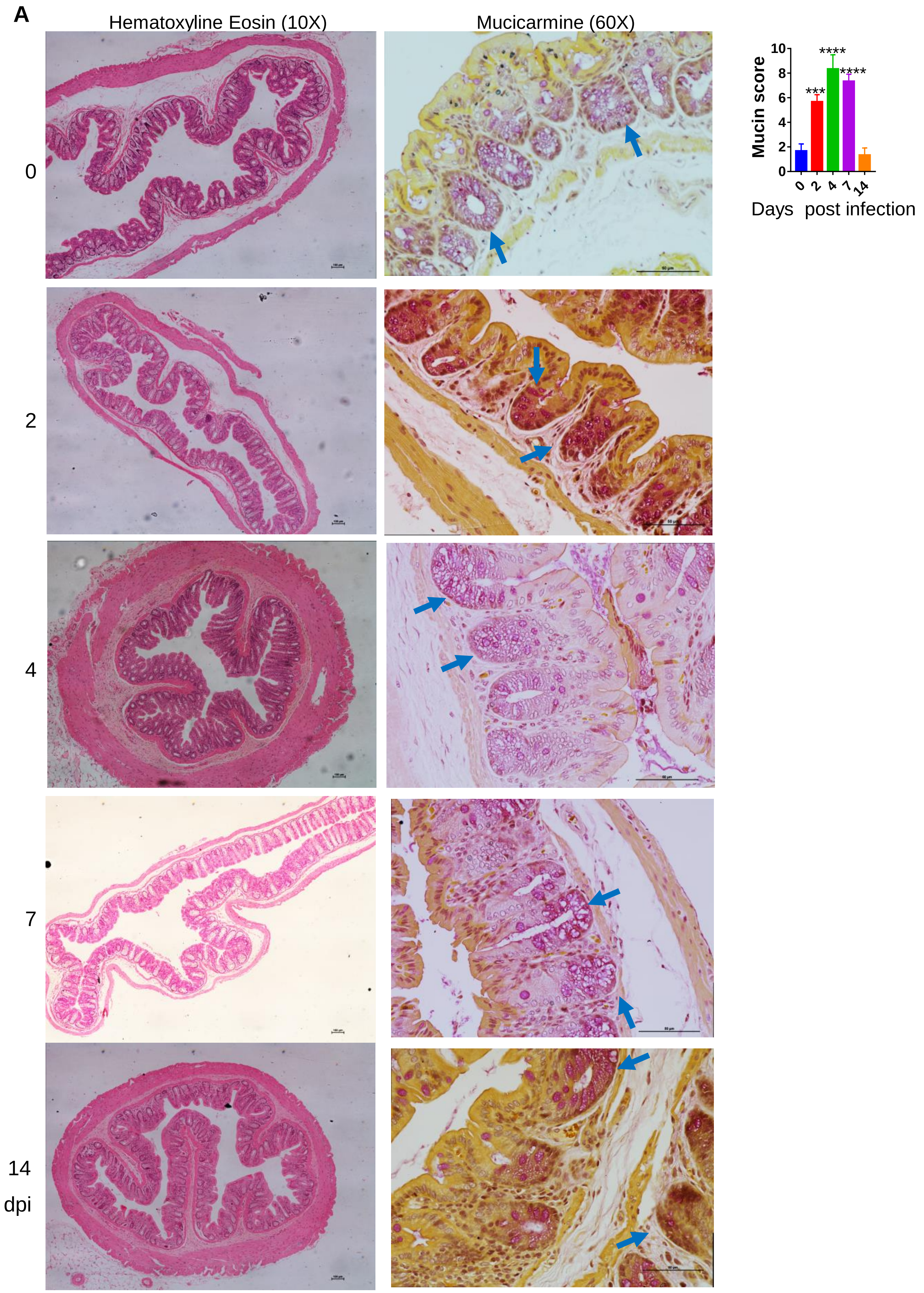

Figure S4

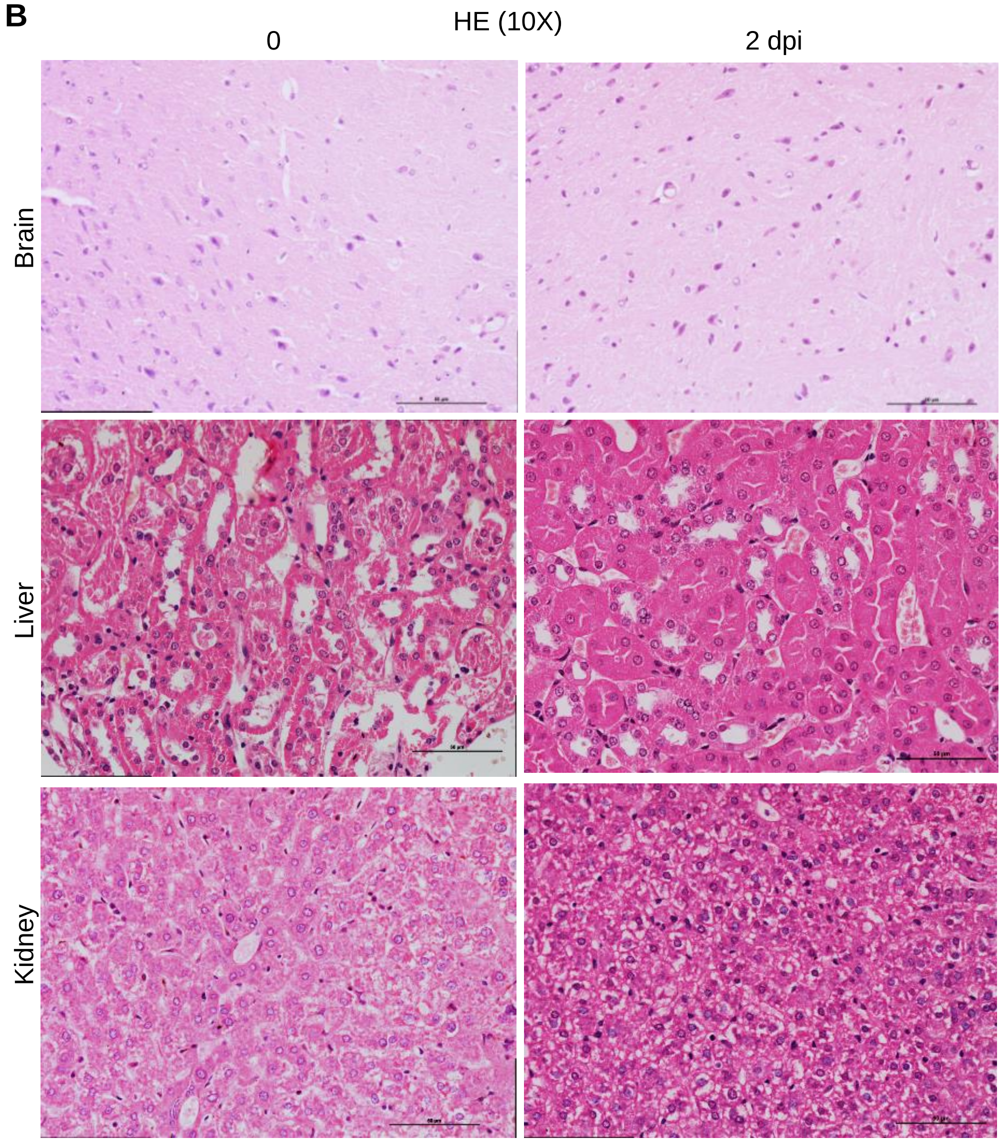

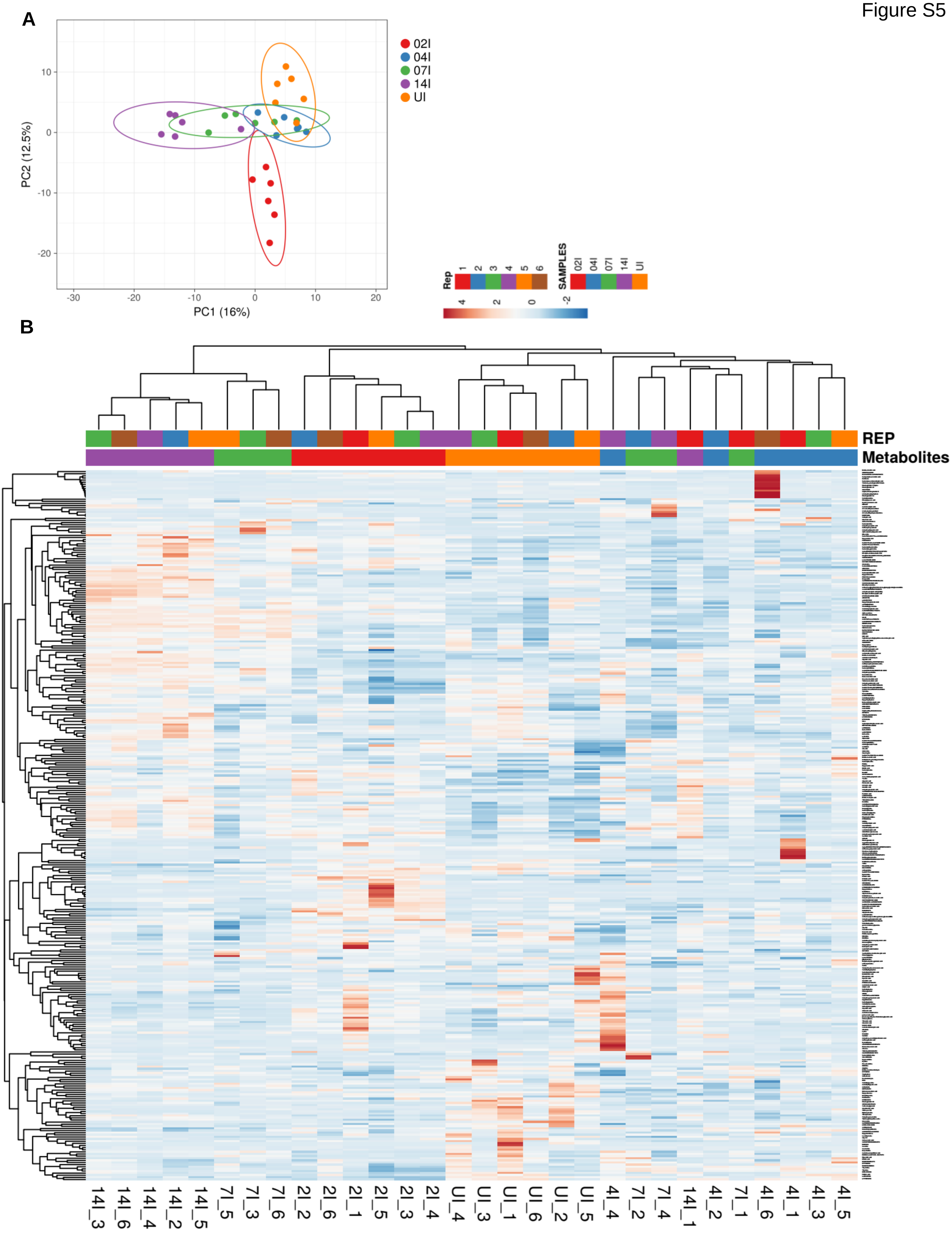

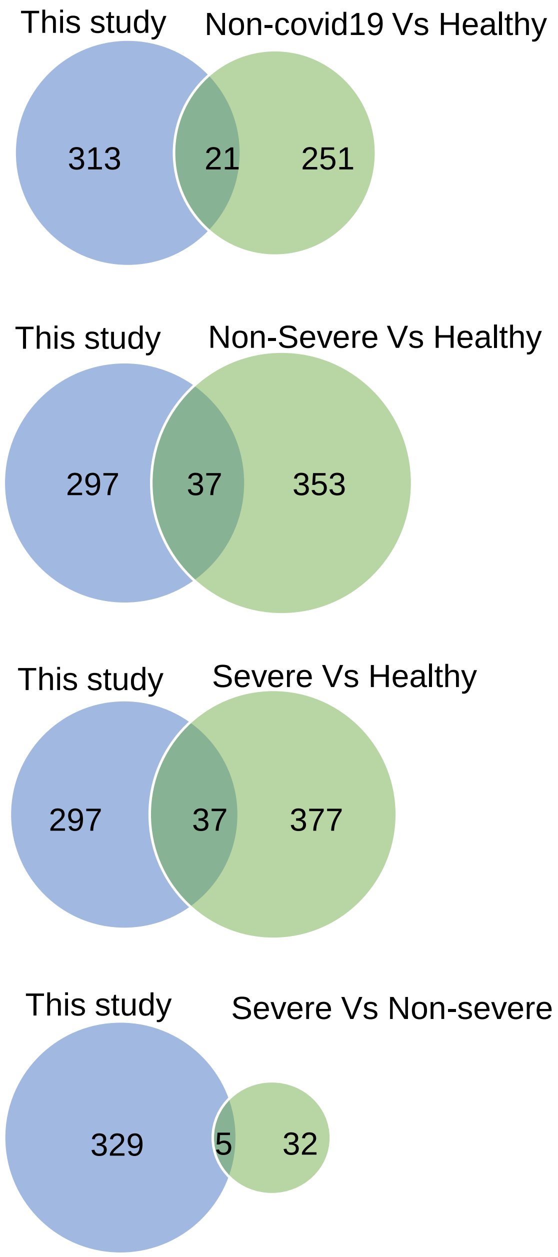

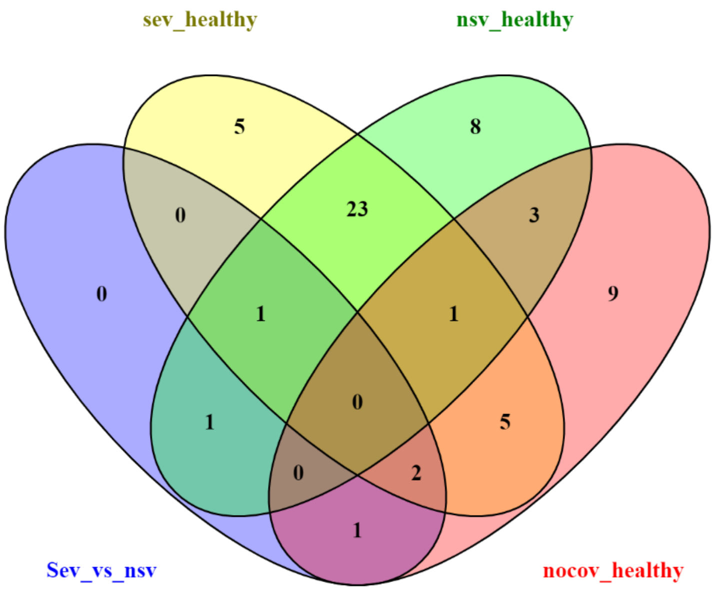
